# Mathematical modelling of thermal death kinetics to describe humane slaughter procedures via hot water immersion in black soldier fly and yellow mealworm larvae

**DOI:** 10.64898/2026.10.14.758861

**Authors:** FS Almeida, MP Zacarias, SO Durosaro, BAM Carciofi, M Barrett

**Author notes:** co-last authors (BAM Carciofi, M Barrett). **Funding:** MB and BAMC received funding from Rethink Priorities to complete this work (in 2025).

## Abstract

Trillions of black soldier fly (BSFL; *Hermetia illucens;* Diptera: Stratiomyidae) and yellow mealworm (YML; *Tenebrio molitor;* Coleoptera: Tenebrionidae) larvae are slaughtered annually for feed. Slaughter standard operating procedures (SOPs) that protect welfare and product quality are essential to ethical and economical industry practice. This study defines humane (≤1 second time-to-death) slaughter SOPs for hot water immersion of BSFL and YML. We assessed isothermal critical thermal maxima in water from 32, or 40, to 100 °C in YML and BSFL, respectively. We used Weibull probability-based models to characterize survival as a function of processing time at different temperatures, describing the time–temperature conditions required for humane slaughter. We blanched larvae under: 1) a SOP resulting in a ∼1 minute time-to-death for 99% of larvae; 2) a SOP resulting in ≤1 second time-to-death for 99% of larvae; and 3) 99.5-100 °C. We compared ash, protein and fat, pH, water activity, lipid oxidation, and color change. Water temperatures kept above 81.3 °C (BSFL ≤ 0.16 g), and 78.3 °C (YML ≤ 0.12 g), are predicted to result in humane slaughter for >99% of larvae - though, further research is needed to develop SOPs for larger larvae. The humane SOPs have neutral or beneficial impacts on quality. We recommend producers adopt a 2 °C buffer (BSFL: 83.3 °C; YML: 80.3 °C). Finally, we provide an energy balance model that producers can use to determine the larvae:water ratio and temperature that results in humane slaughter in their facility.

## 1. Introduction

At least 2-3 trillion black soldier fly (BSFL; *Hermetia illucens*, Diptera: Stratiomyidae) and yellow mealworm (YML; *Tenebrio molitor*, Coleoptera: Tenebrionidae) larvae are farmed and slaughtered each year for food and feed (McKay and Shah, 2025). The number of larvae slaughtered per year is expected to grow by nearly 1000% by 2033 as a result of the insect as food and feed industry’s rapid growth (McKay and Shah, 2025). This novel agriculture industry aims to increase agroeconomic circularity (Lalander *et al*., 2025) through waste reclamation (Rehman *et al*., 2023) and protein production with less land, water, and energy usage than some farmed vertebrates (Hancz *et al*., 2024; Purkayastha and Sarkar, 2022).

Currently, there are no welfare protections for farmed insects, however academics and industry professionals alike have expressed a desire to minimize potential welfare issues for farmed insects even while research on sentience is still ongoing (Andrews *et al*., 2024; Barrett and Fischer, 2023; IPIFF, 2019; Klobučar and Fisher, 2023; van Huis, 2021). Alongside the ethical reasons for caring about insect welfare, animal welfare (especially during slaughter) is often important to the public and can be important for maintaining an animal use industry’s social license to operate (Barrett and Adcock, 2023). Improved insect welfare standards have thus been identified as an important part of the business case for insect farming in Europe (Niyonsaba *et al*., 2024).

To provide humane slaughter for farmed animals, methods would either need to be intrinsically painless, occur with prior stunning to result in unconsciousness during slaughter, or occur in one second or less (i.e., ‘instantaneously’). Little is known about what methods could be intrinsically ‘painless’ for insects if they are sentient (though, plausibly heat, cold, mechanical damage, and at least some chemicals have the potential to activate nociceptors; Goldberg *et al*., 2024; Im and Galko, 2012). Further, no validated stunning methods exist for farmed insects at this time (Barrett *et al*., 2023). Therefore, determining which methods of slaughter have the potential to be ‘instant’ is essential to providing humane slaughter for insects.

Minimal evidence-based guidance exists for producers on the question of what methods or standard operating procedures might result in a humane death for their animals (Bakker *et al*., 2026). Barrett *et al*., (2023b, 2023a) suggest that grinding, boiling/blanching, and freezing in liquid nitrogen have the potential to result in humane (i.e., instantaneous), but that standard operating procedures would need to be further tested for each method. Of these, only humane standard operating procedures for grinding have been developed (Barrett *et al*., 2024; Zacarias *et al*., 2025). Hot water immersion (sometimes colloquially referred to as ‘boiling’ even if the water is below boiling temperature) is a commonly used slaughter method and can be expected to result in significant distress for larvae if death is not instant. For instance, immersion in rapidly heating water causes standard nocifensive behaviors in BSFL, including thrashing their bodies, curling, and contracting, before seizing and paralysis (death) occur (Durosaro and Barrett, 2026).

Although these behaviors suggest that immersion in hot water can be highly noxious to BSFL, as is the case with other insect larvae (Chattopadhyay et al., 2012), if the water temperature is high enough it may be possible for BSFL and YML to die instantly (≤1 second). A favorable surface-area-to-volume ratio and small mass means that heat transfer from water to larvae could be extremely fast. Accordingly, in this study, we set out to define parameters that can lead to humane (≤1 second time-to-death) slaughter for BSFL and YML via hot water immersion (e.g., boiling or blanching). First, we investigated the relationship between constant water temperature and death time (defined via behavioral ‘paralysis’, taken to indicate heat coma onset and loss of sensibility (Jørgensen et al., 2020), generally following seizure), in individual larvae of both species via an isothermal critical thermal maxima assay. Next, we applied Weibull probability-based models to these data to describe the survival likelihood as a function of processing time at each different temperature tested. The goal was to model the fraction of larval survival throughout hot-water immersion and thereby describe the time–temperature conditions required for effective slaughter in 1 second or less.

To assess the impact of humane slaughter on product quality, we chose three different standard operating procedures to be evaluated based on model outputs for each species: 1) an ‘inhumane’ death condition that resulted in ∼1 minute long death times for larvae; 2) a ‘humane’ death condition that resulted in instant death for larvae, at minimized necessary energy use; and 3) an ‘overkill’ condition (99.5 - 100.0 °C) that would also result in instant death for larvae, but with potential increased energy use. Larval flour quality was evaluated by crude protein, crude fat, and ash content, and pH, water activity, and lipid oxidation. Finally, to assist producers in determining the parameters that will provide humane slaughter for their insects, we built an energy balance model that allows producers to input key parameters (larval mass and temperature, water volume, etc.) and determine the water temperature needed to provide instantaneous death for 99% or 99.99% of their animals.

## 2. Material and methods

### 2.1. Animal sourcing

Animals were sourced from Fluker Farms (Port Allen, LA) directly, or from established laboratory populations in the Barrett lab that were originally sourced from Fluker Farms. Black soldier fly larvae were reared on Gainesville diet (Hogsette, 1992); yellow mealworm larvae were reared on 20% nutritional yeast, 80% wheat bran (Dobrowolski *et al*., 2026) with agar gel *ad libitum* as a moisture source. Animals were reared in an incubator (Perceval DR41-VL) with 12:12 L:D, 70% RH, and 27 °C. For further details on standard husbandry practices, see those previously described (Barrett 2026a, Barrett 2026b).

### 2.2. Isothermal critical thermal maxima assays

BSFL that were 0.13 - 0.16 g (mean = 0.14 ± 0.009 g) in body mass were selected as well as YML that were between 0.074 - 0.12 g (mean = 0.11 ± 0.009 g). Animals were briefly cleaned in distilled water, then dried with a paper towel before weighing on an analytical balance to the nearest 0.1 mg (VWR - 220TC). A 50 mL glass beaker was filled with 25 mL of distilled water (‘inner beaker’); heavy magnetic weights (Gorilla Automotive) were superglued (Loctite) to the bottom of the base of the inner beaker, and it was placed inside another 500 mL glass beaker (‘outer beaker’) which was filled with distilled water, such that the water line inside the inner beaker was always underneath the water line of the outer beaker.

An Omega K-type thermocouple (time constant < 0.25 s) attached to an HH506A Omega thermologger was placed inside the distilled water of the inner beaker >1 cm under the surface of the water. As water was deionized, and the immersion depth was greater than 10 times the diameter of the probe, galvanic and thermal conductivity errors of using an exposed probe could be minimized and thereby the time constant delay and increased water depth that would have been necessary for use of a sheathed probe could be avoided.

The water in the inner beaker was then heated until it reached within 0.5 °C of the median temperatures. For BSFL, the measured temperatures (determined after a short pilot experiment) were (median): 41.3, 48.3, 50.1, 52.1, 54.0, 55.7, 57.7, 57.5, 63.4, 68.2, 69.5, 72.5, 81.8, 90.2, and 99.7 °C. For YML, the measured temperatures were (median): 32.1, 40.3, 42.1, 44.6, 47.8, 50.1, 53.0, 55.4, 61.1, 66.0, 71.7, 79.1, 90.2, 99.5°C. In total, we used n = 95 BSFL and n = 106 YML. For each animal, the temperature of the logger was measured at the time the test started, and this was used as the ‘real temperature’ the animal experienced in analyses. In both species, we began with an n = 3 or 5 animals/temperature and increased sample size iteratively based on where models failed to converge and required more data due to high variance.

After the desired temperature was reached, a video camera (HC-V700, Panasonic Full HD, 4 frames/s) facing the water laterally was started; the larva was dropped into the water and watched for shaking and then paralysis. Following paralysis, larvae were removed from the water and placed on the benchtop and observed for motion for ten more seconds; they were then immediately placed into a vial and frozen at -20 °C. If larvae were not paralyzed in three (BSFL) or two (YML) minutes, they were removed from the water, anesthetized with isoflurane, and then placed in a vial, followed by freezing at -20 C for euthanasia. Frozen larvae were dried in the oven at 70 °C for 12 hours and re-weighed for dry mass.

Videos were watched back and scored for the second that seizure and paralysis started in each species; paralysis following seizure is a commonly-used measure of heat-associated death in insects, including larvae generally and BSFL specifically (Chattopadhyay *et al*., 2012; Durosaro and Barrett, 2026). All animals that were paralyzed in the water failed to walk or crawl when removed from the water; all animals that failed to be paralyzed in the water walked following removal from the water.

Considering that time-to-death was determined by behaviors scored in video recordings, the measurement resolution was limited to 1 s. Below this value, resolution was insufficient to reliably distinguish the time of death.

### 2.3. Energy balance of the larva slaughtering process

The slaughtering of insects in hot water immersion is assumed to be under constant pressure and total mass, without involving phase change as this can be neglected. In this case, the enthalpy variation (Δ*H*) of any part of the system is estimated according to Eq. 1.

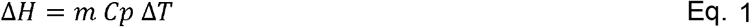

in which *m* is the mass of system (insects, water) (kg), *Cp* is its specific heat capacity (J⋅ kg^-1^ ⋅°C^-1^), and *T* is its temperature (°C).

Figure 1 illustrates the mass and energy flows in a generic system for animal slaughter by immersion in hot water, assuming perfect mixing of the larvae and water.

**Figure 1.**
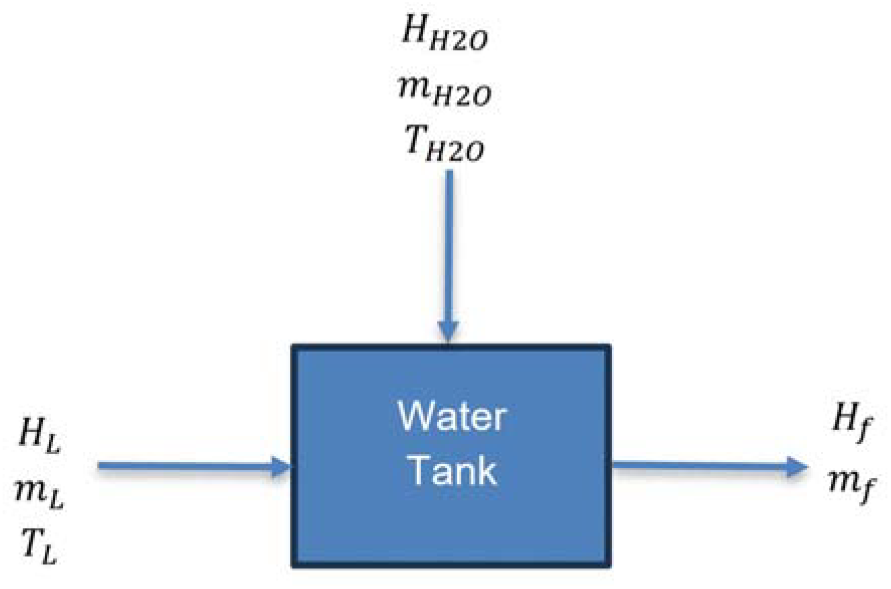
Energy and mass flows for the mixing of larva and water during slaughtering by hot water immersion. The subscripts represent: *L* for larva properties; *H*2*O* for water properties, and *f* for the final system (perfect mix of larva and water).

By mass conservation, the final mass (*m*_*f*_) is the sum of the larva mass (*m*_*L*_) and the water incorporated into the tank (*m*_*H*2*O*_) (Figure 1).

According to the diagram in Figure 1, the energy balance neglecting any energy losses (to the environment or by phase change) and mixing enthalpy will result in the Eq. 2.

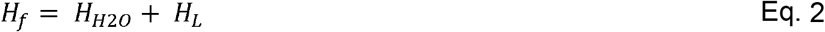

in which *H*_*f*_ is the final enthalpy of water-larvae mixture at equilibrium (assuming perfect mixing), *H*_*H*2*O*_ is the initial enthalpy of the water added to the tank, and *H*_*L*_ is the enthalpy of the incoming larvae.

By combining Eq. 1 and Eq. 2, it is then possible to estimate key parameters of the slaughtering process by isolating the desirable output. Different scenarios can be considered such as: i) the estimate of the the system (larva+water) final temperature, *T*_*f*_, for a given initial water temperature (*T*_*H*2*O*_), larva mass (*m*_*L*_), and water mass (*m*_*H*2*O*_) (Eq. 3); ii) the estimate of the water mass, *m*_*H*2*O*_, to be added targeting to reach a certain final temperature when using a known mass of larvae and initial water temperature (Eq. 4); iii) the estimate of the larval biomass, *m*_*f*_, allowed to be added to a given water quantity at a certain temperature to achieve a final desirable temperature (Eq. 5). A spreadsheet with this model implemented has been added to the Supplementary Material (Supplemental File 1) to assist in determining the process parameters that will provide humane slaughter for the insects, while allowing the addition of input key parameters.

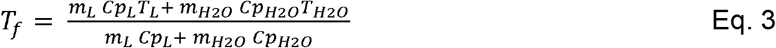

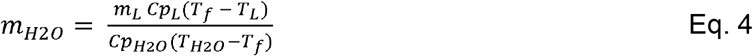

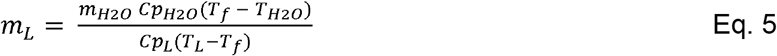

in which *C*_*H*2*O*_ and *C*_*pL*_ are the specific heat capacity of water (4.187 J ⋅ kg^-1^ ⋅ °C^-1^) and larvae (estimated with Eq. 6), respectively. In estimates presented in the Supplementary Material (Supplemental File 1), the initial larvae temperature was assumed to 21 °C. Heldman and Singh (1981) proposed that the specific heat capacity of biological materials can be estimated as an additive property based on their proximate composition (Eq. 6).

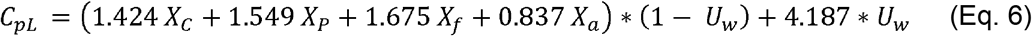

Where *X* is the mass fraction (g component/g dry solids) and the subscripts represent carbohydrate (*C*), protein (*P*), fat (*f*), and ash (*a*). *U*_*w*_ represents the moisture content on a wet basis (g water/g total solids).

The literature reports varying data on the proximate composition of BSFL and YML (Supplemental File 2) which, for crude protein, may be in part due to the use of different protein to nitrogen conversion factors (SUSINCHAIN, 2023). We therefore calculated the average mass fraction of each component from the literature to determine the specific heat capacity of each insect species (Supplemental File 3).

### 2.4. Mathematical model of YML and BSFL time-to-death, death kinetics, and probability of survival as a function of water temperature

For model development, we began by grouping the observed time-to-death values for individual larvae according to the reference temperature to which they were exposed. In doing that, we excluded any data points that were more than 1 °C away from the set temperature, except in the case where that would cause overlap between two adjacent set temperatures. In that case, for temperature ranges in which the time-to-death was more sensitive to temperature variations, we excluded any data points that were more than 0.5 °C above or below the set temperatures. Thus, a total of 8 data points were removed from the modeling dataset for BSFL and none for YML. We also did not model temperatures where the animals did not die before the cut off time. Altogether, 88 datapoints for BSFL and 74 for YML were used to fit and elaborate the models.

Aiming at predicting the survival rate or probability of survival of larvae through treatment time after undergoing slaughtering in hot water immersion, a multi-step modeling approach was proposed (Figure 2). First, time-to-death data grouped at each reference temperature (isothermal condition) was fitted by the Weibull probability distribution for each larva (BSFL and YML) (Eq. 7). The resulting probability density function was then integrated to obtain the cumulative distribution function (CDF, Eq. 8). Finally, the survival fraction was calculated as the complement of the CDF (1 – CDF, Eq. 9).

**Figure 2.**
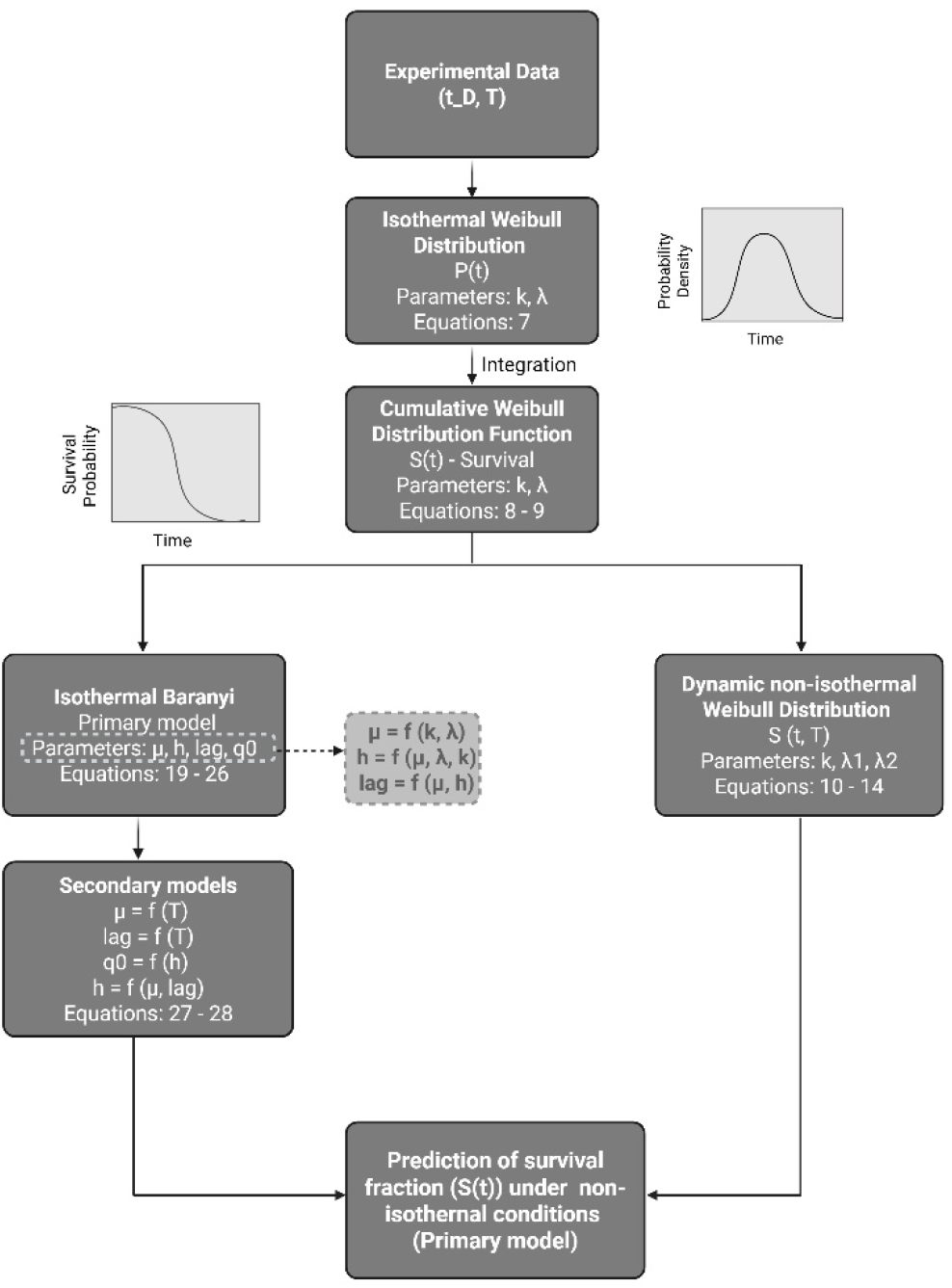
Modeling approach used to fit probability distributions to experimental data (temperature and time-to-death).

Although the survival fraction describes the probability of larval survival over time, it does not explicitly account for the cumulative thermal injury experienced by the larvae under non-isothermal conditions during slaughtering. Therefore, temperature variation and cumulative thermal injury were then incorporated into the Weibull model to describe larval survival under non-isothermal conditions, resulting in the development of the dynamic Weibull distribution (Eq. 10 – 14). Likewise, a Weibull–Baranyi approach was also evaluated, as it accounts for both the effective kinetic parameter and the cumulative stress associated with thermal injury. When fitted to isothermal conditions (Eq. 19 – 26), the model parameters can be further used to develop secondary models (Eq. 27 – 28), enabling the estimation of survival probabilities at temperatures not experimentally evaluated but within the studied temperature range. These selected models present a sigmoidal shape and are often used to describe the kinetics of microorganisms (Ranjbaran *et al*., 2021). A similar approach translating sigmoid model to insect’s survival was previously proposed by Parisotto *et al*., 2023.

#### Isothermal survival model based on Weibull distribution

The Weibull distribution was used to describe the distribution of time-to-death under isothermal conditions. This two-parameter distribution is widely applied in reliability and survival analyses because of its flexibility in representing different survival behaviors. The shape parameter (*k*) determines the form of the distribution, whereas the scale parameter (*λ*) defines its scale (Hodges, 1994; Weibull, 1951).

The probability density function (PDF) of the Weibull distribution is expressed as (Eq. 7):

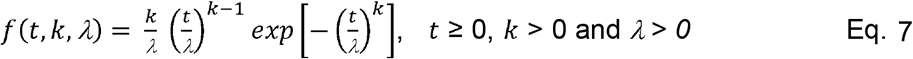

where *t* is the treatment time, *k* is the shape parameter, and *λ* is the scale parameter.

The corresponding integrative or cumulative distribution function (CDF), which represents the probability that thermal death has occurred by time *t*, is given by Eq. 8:

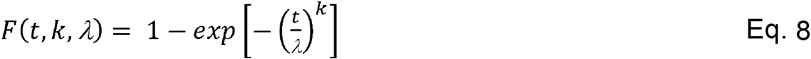

Consequently, the survival function, representing the probability of an individual remaining alive after time, was calculated as (Eq. 9):

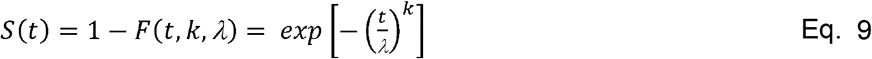

The Weibull distribution was selected because it is restricted to positive times (*t* ≥ 0), making it appropriate for modeling time-to-death data. Other distributions such as the normal distribution may present negative treatment times (*t* < 0), which have no physical meaning in thermal inactivation experiments. The distribution was generated in the interval average time-to-death ± 4 standard deviations, and with 1000 points in between this range.

#### Dynamic Weibull Distribution Shape parameter as function of temperature (λ T))

By introducing the shape parameter *λ* (*T*) as a function of temperature, the survival function can be rewritten as (Eq. 10):

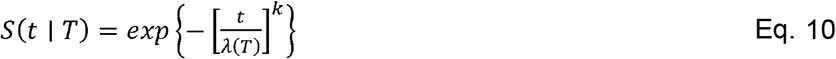

And *λ* (*T*) can be described by the following function (Eq. 11):

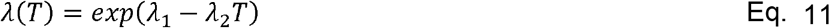

where *S*(*t,T*) is the survival probability at exposure time *t* and constant temperature *T, k* is the common Weibull shape parameter, *λ*(*T*) is the scale parameter as a function of temperature, and *λ*_1_ and *λ*_2_ are fitting parameters.

#### Cumulative thermal injury (Q)

To describe larval survival under non-isothermal conditions, the Weibull survival function was further modified by incorporating a cumulative thermal injury variable *Q*(*t*). This variable, often used in microbial thermal inactivation models, represents an organism’s susceptibility to its environment as a function of its prior thermal history (Longhi *et al*., 2018; Ranjbaran *et al*., 2021). Translating that to the present case, *Q*(*t*) introduces into the equation the effect of the fluctuating temperatures experienced by the larva over time, accounting for the progressive physiological damage such as protein denaturation and loss of cellular function, which accumulates as the animal is exposed to sub-lethal thermal conditions, ultimately influencing its probability of survival.

Under non-isothermal exposure, the temperature dependence enters through the rate of injury accumulation (Eq. 12 and 13):

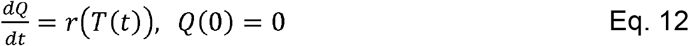

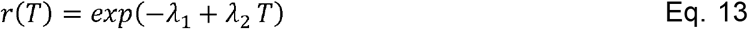

The survival probability *S*(*t*) is then obtained from *Q*(*t*) as (Eq. 14):

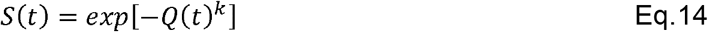

By solving simultaneously Eq. 12, 13 and 14 and fitting all the isothermal data together, we can then obtain the numerical simulation of non-isothermal survival (Dynamic Weibull Distribution).

#### Weibull-Baranyi survival model

This approach relies on a dynamic ordinary differential equation (ODE)-based modeling that combines a Weibull cumulative distribution function with a Baranyi-type structure (Baranyi and Roberts, 1994), suitable for non-isothermal conditions. In this case, we target the development of a primary model that describes the thermal death of YML and BSFL under isothermal conditions and subsequently derive a secondary model from the primary model’s parameters. By combining both models, this strategy allows the modelling of larval thermal death at temperatures not directly tested within the evaluated range, since the primary model’s parameters can be estimated from the secondary model, enabling reconstruction of the corresponding survival probability curves.

#### Primary model

For this, the modelling can be described in terms of mortality accumulation (*D*(*t,T*))) or cumulative fraction of dead individuals (Eq. 15), which is derived from Eq. 9.

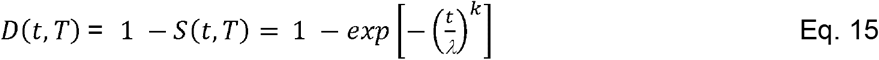

Moreover, a dynamic Baranyi-type differential system is proposed (Eq. 16 – 18), with the following initial conditions: *D*(0) = ε *q*(0) = *q*_0_ (Ranjbaran *et al*., 2021).

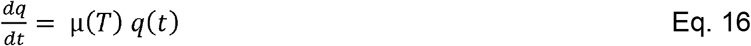

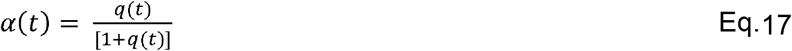

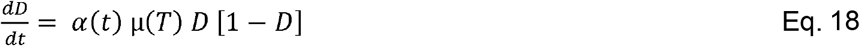

where μ(*T*) is the effective kinetic parameter of the Baranyi-type model, *q*(*t*) is the cumulative stress due to thermal injury and *ε* is a small positive number introduced (0<*ε* ≪ 1) to avoid the singularity associated with starting at *D* = 0.

By solving together Eq. 16 to 18, it is then possible to estimate *k*(*t*) and *λ*(*t*), from the observed death-time data, and obtain values for μ(*T*) (death rate) and *q*_0_ (initial physiological state parameter). Consequently, the resulting differential system can then be solved for non-isothermal histories *T* = *T*(*t*).

Fittings were performed under isothermal conditions using the following integrated equations (Eq. 19 – 20) (Laroque *et al*., 2023). Moreover, estimates of μ, *q*_0_, *lag* and *h* can be derived from Eq. 23 – 26, using the shape (*k*) and scale (*λ*) parameters from the Weibull distribution.

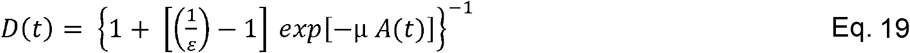

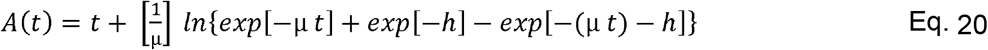

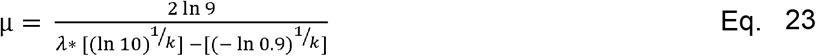

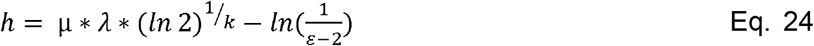

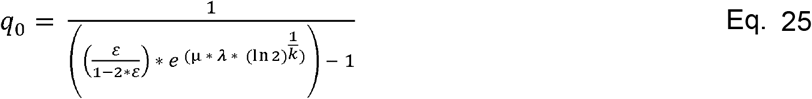

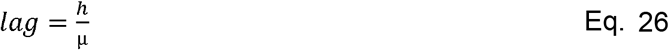

Where *lag* represents the adaptation period of the population before the main inactivation process becomes fully expressed.

#### Secondary model

Secondary models were employed to describe the temperature dependence of the primary model parameters (μ and *lag*). These models enable the prediction of survival probability curves for YML and BSFL under isothermal conditions that were not directly investigated experimentally.

Two secondary models were developed: a linear model to describe the relationship between the kinetic parameter (μ) and temperature (*T*) (Eq. 27), and a second hyperbolic model to characterize the nonlinear relationship between the parameter (*lag*) and temperature (*T*) (Eq. 28). Given the inherent variability of biological data, data points that deviated substantially from the overall observed trends were excluded from the model fitting.

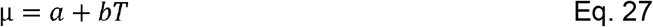

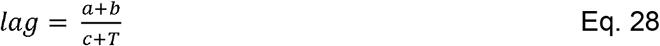

### 2.5. Product quality assays

For BSFL, 145 g (Bonvoisin HZ30002B balance) of live larvae were weighed per sample (∼1,950 larvae, mean individual mass of 0.074 g; VWR220-TC). Each BSFL sample was then randomly assigned to one of three treatment conditions: 1) ‘inhumane’ death condition: 52.0 °C for 55 seconds; 2) humane’ death condition: 83.3 °C for 1 second; or 3) ‘overkill’ condition: 99.5 to 100.0 °C for 1 second. For YML, 145 g of live larvae were weighed per sample (∼1,750 larvae, mean individual mass of 0.083 g). Each YML sample was then randomly assigned to one of three treatment conditions: 1) 47.0 °C for 55 seconds; 2) 80.3 °C for 1 second; or 3) 99.5 to 100.0 °C for 1 second.

For both species, 8 L of water was heated and the temperature maintained by an Inkbird ISV-100W sous vide (for procedures 1 and 2) or by boiling the water in an aluminum cooking pot on a Duxtop 1800 W induction heater. K-type stainless-sheathed thermocouples (PerfectPrime TL1915SL) were immersed in the water and attached to Omega HH506A thermologgers to confirm water temperature. Immediately prior to the test, the larvae were placed in a mesh spaghetti-cooking basket, with a second basket placed over top to prevent floating. The basket was then immersed in the water for the requisite amount of time. Following immersion, larvae were immediately removed from the basket and placed into a cooler, and ∼2 L of liquid nitrogen was poured over the sample, which was stirred with metal tweezers. Excess liquid nitrogen was poured off, and the sample was transferred in a plastic bag to a -80 °C freezer.

Samples were briefly crushed with a chilled metal hammer, lyophilized (BSFL: freezing for 61 min, vacuum freezing for 27 min to -24 °C and 59 Pa, primary drying for 15 h 50 min with a temperature ramp from -40.0 °C to -51.7 °C, and final drying drying at -51.7 °C under 22 Pa vacuum for 3 h. YML: freezing for 60 min, vacuum freezing for 17 min to -31 °C and 60 Pa, primary drying for 10 hr 24 min with a temperature ramp from -40.0 °C to -51.7 °C under 22 Pa vacuum for 4 h 20 min. Both BSFL and YML were lyophilized with a HarvestRight Freezer Dryer, Scientific Model), then ground with a chilled mortar and pestle, and weighed to get dry mass; this was considered the ‘flour’. Larval flour samples were stored in a -20 °C freezer until all samples had been lyophilized, at which point they were sent off to Eurofins Nutritional Analysis for analysis. Eurofins completed ash (AOAC 942.05), crude fat by acid hydrolysis (AOAC 954.02), crude protein (combustion - AOAC 990.03; AOAC 992.15), pH (AOAC 981.12), water activity (AOAC 978.18), and thiobarbituric acid rancidity (Tarladgis et al. 1960) testing; n = 3 samples were used per protocol, per product quality measure. For crude protein, we used a nitrogen to protein conversion factor of 4.75 for *T. molitor* and 4.67 for *H. illucens* (instead of 6.25, but see Supplemental Tables 1 and 2 for values using 6.25) as is recommended for insect protein (Janssen *et al*., 2017; SUSINCHAIN, 2023).

**Table 1.**
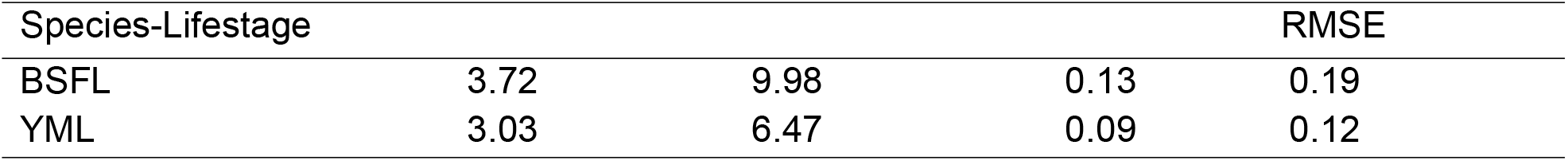
Fitting parameters for the Weibull survival model incorporating a cumulative thermal injury variable.

**Table 2.**
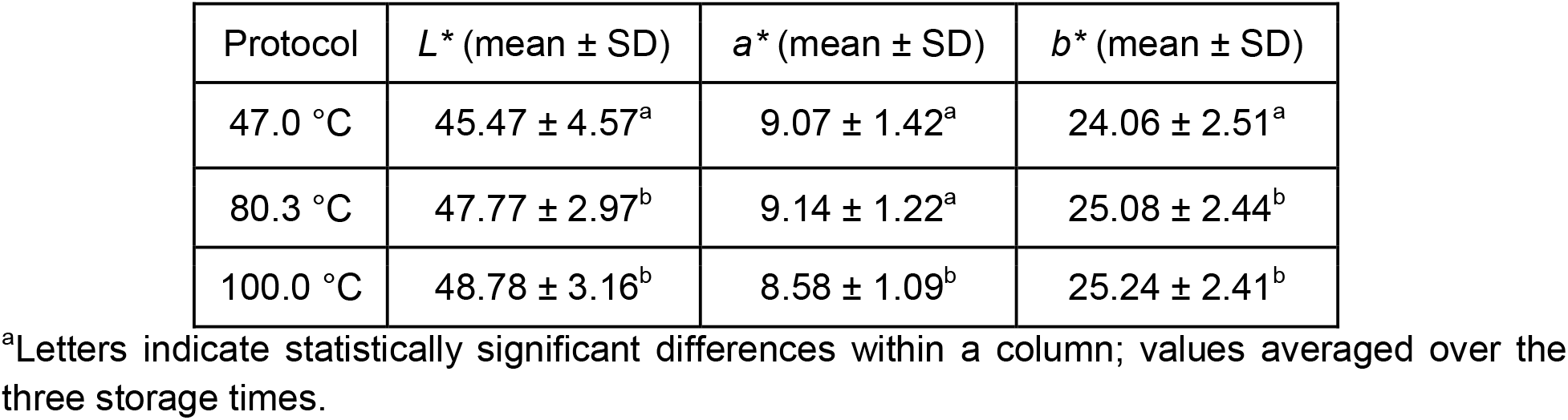
YML flour characteristics (L^**\***^, **a**^**\***^, **b**^**\***^) across protocols.

| Protocol | $L^*$ (mean ± SD) | $a^*$ (mean ± SD) | $b^*$ (mean ± SD) |
| --- | --- | --- | --- |
| 47.0 °C | 45.47 ± 4.57<sup>a</sup> | 9.07 ± 1.42<sup>a</sup> | 24.06 ± 2.51<sup>a</sup> |
| 80.3 °C | 47.77 ± 2.97<sup>b</sup> | 9.14 ± 1.22<sup>a</sup> | 25.08 ± 2.44<sup>b</sup> |
| 100.0 °C | 48.78 ± 3.16<sup>b</sup> | 8.58 ± 1.09<sup>b</sup> | 25.24 ± 2.41<sup>b</sup> |

<sup>a</sup>Letters indicate statistically significant differences within a column; values averaged over the three storage times.

### 2.6. Color analysis

Samples of YML and BSFL flour prepared as in the ‘product quality assays’ section above. 5 g each of either YML or BSFL flour were then placed in a petri dish (diameter: 5.5 cm, height: 1.2 cm) and photographed in a light box (39 cm LWH, Neewer) from a dorsal orientation using a Google Pixel 6a phone on days 0 (the day they were removed from the lyophilizer and ground), 7 and 30. Storage of the flour was in the dark at room temperature (23.97 ± 0.45 °C). Photos were analyzed in ImageJ (Schneider et al. 2012) by converting them to a lab stack and using a 1.8 cm diameter circle to collect mean *L*^*\**^, *a*^*\**^, and *b*^*\**^ values from each photo in three places on the sample (avoiding any shadows); *L*^*\**^, *a*^*\**^, and *b*^*\**^ values were also collected from a white standard in each photo and used for offset corrections. The average, offset-adjusted *L*^*\**^, *a*^*\**^, and *B*^*\**^ values from each photo were used in the final analysis. Photos were taken of the same samples on day 0 (the day of removal from the lyophilizer), day 7, and day 30. For YML, 13 batches per protocol were used (n = 39). For BSFL, we had fewer batches available due to lower flour volumes produced from each batch; therefore, only two batches per protocol with three technical replicate flour samples per batch/protocol combination were used (n = 18) and we accounted for batch identity in subsequent statistics.

### 2.7. Statistical analyses and goodness of fit

Statistical analyses were run in GraphPad Prism (v11.0.1) and R (v4.4.2). Data were tested for normality (Shapiro-Wilk) and heteroskedasticity prior to choosing an analysis method. Alpha was set to 0.05. The effects of protocol on pH and rancidity were assessed using a Kruskal-Wallis with Dunn’s multiple comparisons test post-hoc (for non-normal data) and an ANOVA with Holm-Sidak’s MCT, respectively. The effects of slaughtering protocol on ash (%), crude fat by acid hydrolysis (%), and crude protein (%) were assessed using a beta regression with a likelihood ratio test (LRT) to assess if protocol was a significant predictor (given that data are bounded between 0 and 1). If protocol was found to be significant using the LRT, we used the emmeans (Lenth, 2024) package to conduct a Sidak’s post-hoc multiple comparisons tests among protocols. Offset-corrected *L*^*\**^, *a*^*\**^, and *b*^*\**^ values were analyzed separately using linear mixed-effects models (lme4; Bates et al. 2015) in R, with blanching protocol and day (0, 7, 30) as fixed effects, and random intercepts for photo day, replicate, and replicate nested within batch in BSFL to account for repeated measures; in YML, the same fixed effects and random intercepts for photo day and batch were used to account for repeated measures. Fixed effects were tested with F-tests (lmerTest; Kuznetsova et al. 2017), with Tukey-adjusted comparisons of estimated marginal means for day/protocol differences (emmeans; Lenth 2024). Model fittings were performed in Python (Weibull distribution vs. Dynamic Weibull Distribution) and excel (Weibull distribution vs. Baranyi-Weibull), and the coefficient of determination (R^2^, Eq. 28) and root mean square error (RMSE, Eq. 29) were used to evaluate the goodness of each model fitting.

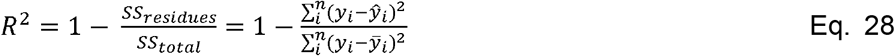

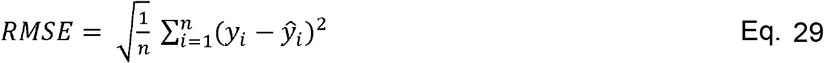

Where *λ*_*i*_ is the actual experimental values, 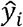 is the predicted or fitted values, 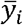 is the global mean, and n is the total number of experimental points.

### 2.8. Ethical use of insects in research statement

Insect research is not subject to welfare legislation in the United States; nevertheless, we sought to consider the 3Rs (Replacement, Reduction, and Refinement) during this work (Fischer *et al*., 2024). For the work with live animals conducted at Indiana University, we developed a voluntary IACUC protocol and received feedback from a volunteer on the IUI IACUC as well as an external reviewer with 10+ years of IACUC experience at another institution. We confirmed lack of duplication or alternative models prior to beginning live animal work, and our choice to use modeling approaches was designed to minimize the number of live animals that would be needed to answer our research questions. We piloted our protocols for the isothermal assay before subjecting a larger number of animals to testing to determine if we needed to study a larger range of body sizes. We further reduced animal use by using an iterative approach to building the model, allowing us to determine how many more data points were needed with each species at each temperature in the isothermal assays, targeting only temperatures that needed more data with new individuals. We used wet-to-dry mass conversion data from our lab population of each species and the optimal sample quantity information provided by Eurofins to estimate how many animals were needed for the product quality tests.

For refinement, we used 3 minutes (BSFL) then 2 minutes (YML, further refined) as our cut-off time for isothermal assays and euthanized any animals alive after this time by isoflurane anesthesia at a saturation dose, followed by freezing at -20 °C. As the behavior and physiology of live, unanesthetized animals undergoing slaughter was an important part of our test, we did not offer any further pain relief or anesthesia to animals in the isothermal or product quality tests. All animals for product quality tests were used within 24 hours of receipt from the supplier (and were shipped without food or water). Food and water were not provided during this time to avoid altering product quality analyses from first samples tested to final samples tested based on feed ingredients.

## 3. Results

### 3.1. Isothermal critical thermal maxima assays

Our pilot tests at three temperatures (BSFL: targeting 56, 64, 72 °C, n = 10 individuals/temp; YML: 44.5, 48, and 79 °C; n = 10 individuals/temp) sometimes showed an effect of large differences in body mass on time to death (see Supplemental Figures 1 and 2); therefore, we preceded using a standardized mass range for each species; we chose a mass range that was slaughter-weight relevant.

0.13 - 0.16 g BSFL (n = 11 tested from 39.8 - 48.9 °C) did not die within 3 minutes at temperatures less than 48.9 °C, as represented by the red area zones in Figure 3. We therefore concentrated our efforts at temperatures from 49.9 - 100 °C (n = 84). 100% of the larvae in our sample (n = 9) tested at temperatures above 81.8 °C died in ≤ 1 second, with the next highest temperature tested at 71.9 - 72.8 °C (and resulting in a 2 - 3 second time to death). 100% of 0.07 - 0.12 g YML (n = 9 tested from 31.8 - 40.5 °C) did not die within 2 minutes at temperatures less than 40.5 °C (Figure 3). Additionally, some individuals did not die within two minutes up to 44.2 °C (n = 2 of 20 individuals tested from 41.7 - 44.2 °C). We therefore concentrated our efforts at temperatures from 44.2 - 100 °C (n = 80). 100% of the larvae in our sample (n = 6) tested at temperatures above 89.7 °C died in ≤ 1 s, with the next highest temperature tested at 79.1 - 79.2 °C and resulting in a 1 or 2 second time to death.

**Figure 3.**
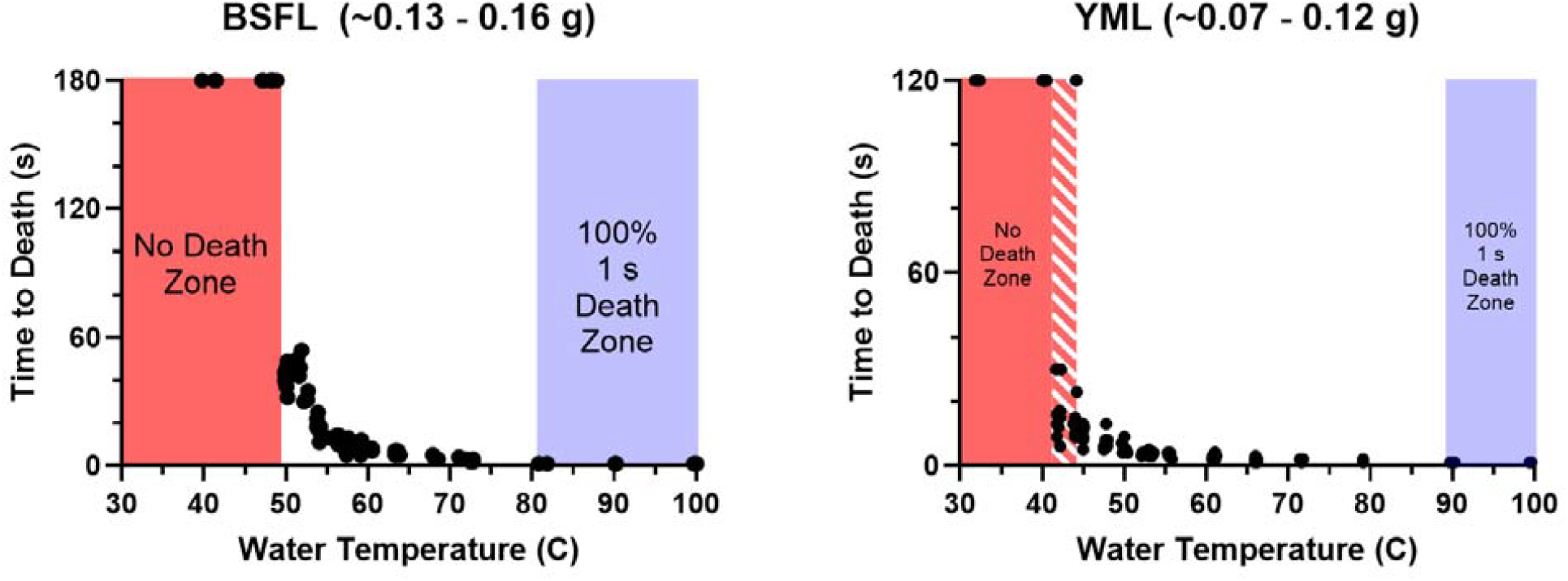
Time to death of individual BSFL and YML tested under isothermal conditions in hot water. Red areas indicate zones where all tested individuals (n = 95 BSFL, n = 106 YML) did not die within the evaluation window (three minutes (BSFL) or two minutes (YML)). Red diagonal hashing indicates temperatures where some, but not all, animals died within the evaluation window. Blue areas indicate tested temperatures where 100% of sampled larvae died in ≤ 1 second (‘instantly’). Note that the points showing as ‘180 s’ for the BSFL and ‘120 s’ for YML are not death times -these are animals that did not die before the cut-off times were reached.

### 3.2. Modelling the probability of insect survival using different modelling approaches

Figures 4 and 5 illustrate the survival probability for BSFL and YML larvae, respectively, as function of the processing time at different temperatures, with the fitted models derived from Weibull statistical probability distributions. Fittings were performed for isothermal treatments ranging from 50 to 72 °C for BSFL and 42 to 66 °C for YML. Temperatures above these ranges, which produced a time-to-death of 1 s and below, were not modeled because the measurement resolution was insufficient to reliably distinguish values below this threshold (≤1 s). Temperatures below these ranges were not modeled as they did not produce death within 3 min (BSFL) or 2 min (YML), our experimental cut-off times.

**Figure 4.**
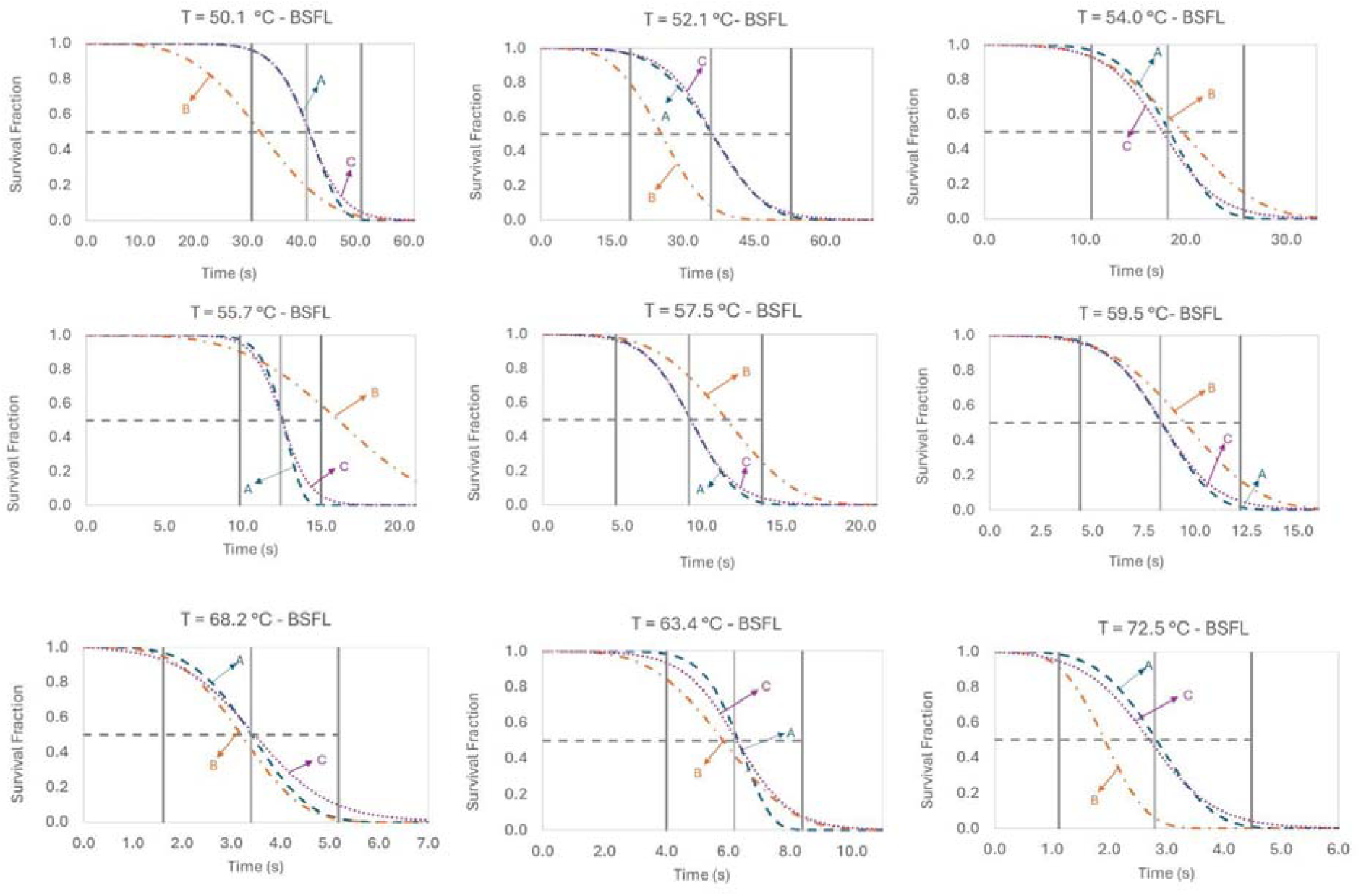
Statistical-based modelling of BSF larvae using Weibull probability distributions. Survival curves over processing time, modeled for temperatures ranging from 50 to 72 °C. Temperatures tested above these thresholds were not modeled as they yielded ≤1 s time-to-death, the measurement resolution for these experiments. The blue curve represents the traditional Weibull distribution (A), the orange curve represents the dynamic Weibull model (B), and the purple curve is the model fitting for the Baranyi distribution (C). The gray dashed horizontal line represents 50% survival, while the filled vertical lines represent the interval between the averaged time-to-death for each temperature ± 2 standard deviations (dark gray lines). Graphs are presented in different time scales to facilitate visualization of the different models.

**Figure 5.**
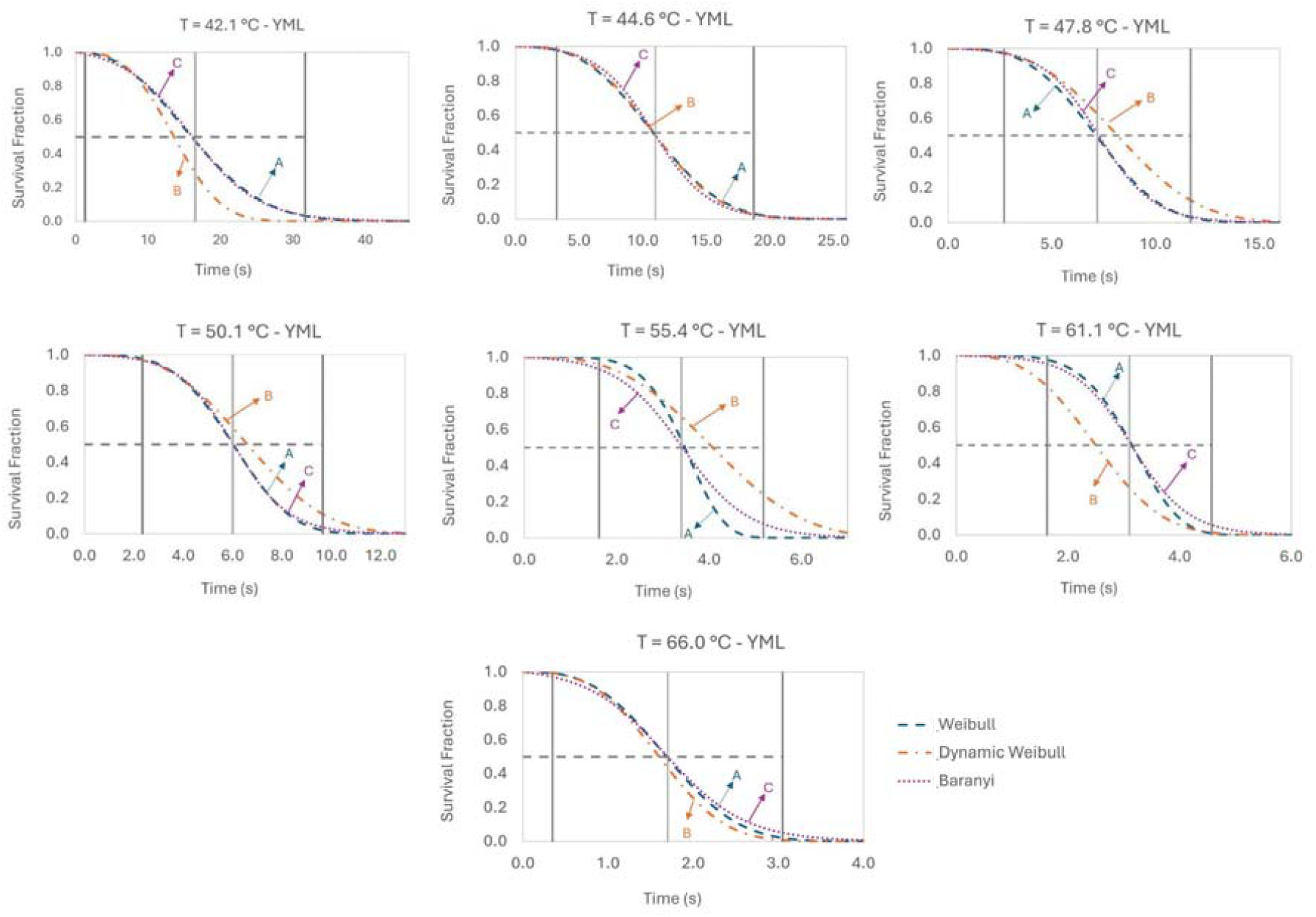
Statistical-based modelling of YML larvae using Weibull probability distributions. Survival curves over processing time, modeled for temperatures ranging from 42 to 66 °C for YML. Temperatures tested above these thresholds were not modeled as they yielded ≤1 s time-to-death, the measurement resolution for these experiments. The blue curve represents the traditional Weibull distribution (A), the orange curve represents the dynamic Weibull distribution (B), and the purple curve is the model fitting for the Baranyi distribution (C). The gray dashed horizontal line represents 50% survival, while the filled vertical lines represent the interval between the averaged time-to-death for each temperature ± 2 standard deviations (gray lines). Graphs are presented in different time scales to facilitate visualization of the different models.

The average time-to-death, defined as the time required to reach 50% survival (*S*= 0.5), was lower for YML than for BSFL under the same isothermal conditions at the lower to middle end of the tested temperature range. For example, at 50.1 °C, the average time-to-death was 6.0 s for YML compared with 41.0 s for BSFL, respectively. Increasing temperature decreased the time required to reach the inflection point of the survival curves, resulting in shorter processing times for the slaughter of both BSFL and YML. From the isothermal data (Figures 4 and 5), the Baranyi survival model was observed to closely follow the Weibull distribution’s behavior for both species, often presenting the lowest RMSE values for each isothermal condition (Supplemental Material, Supplemental Table 3). This is likely because the Baranyi model’s form is similar to a sigmoidal shape and relies on the shape (*k*) and scale (*λ*) parameters from the traditional Weibull distribution, as shown in Eq. 19 – 26. Additionally, this distribution shows that at 50% survival probability (*S*= 0.5), the time-to-death model closely matches the average time-to-death observed (e.g., indicated by the middle gray vertical line in Figures 4 and 5) for each evaluated temperature.

**Table 3.**
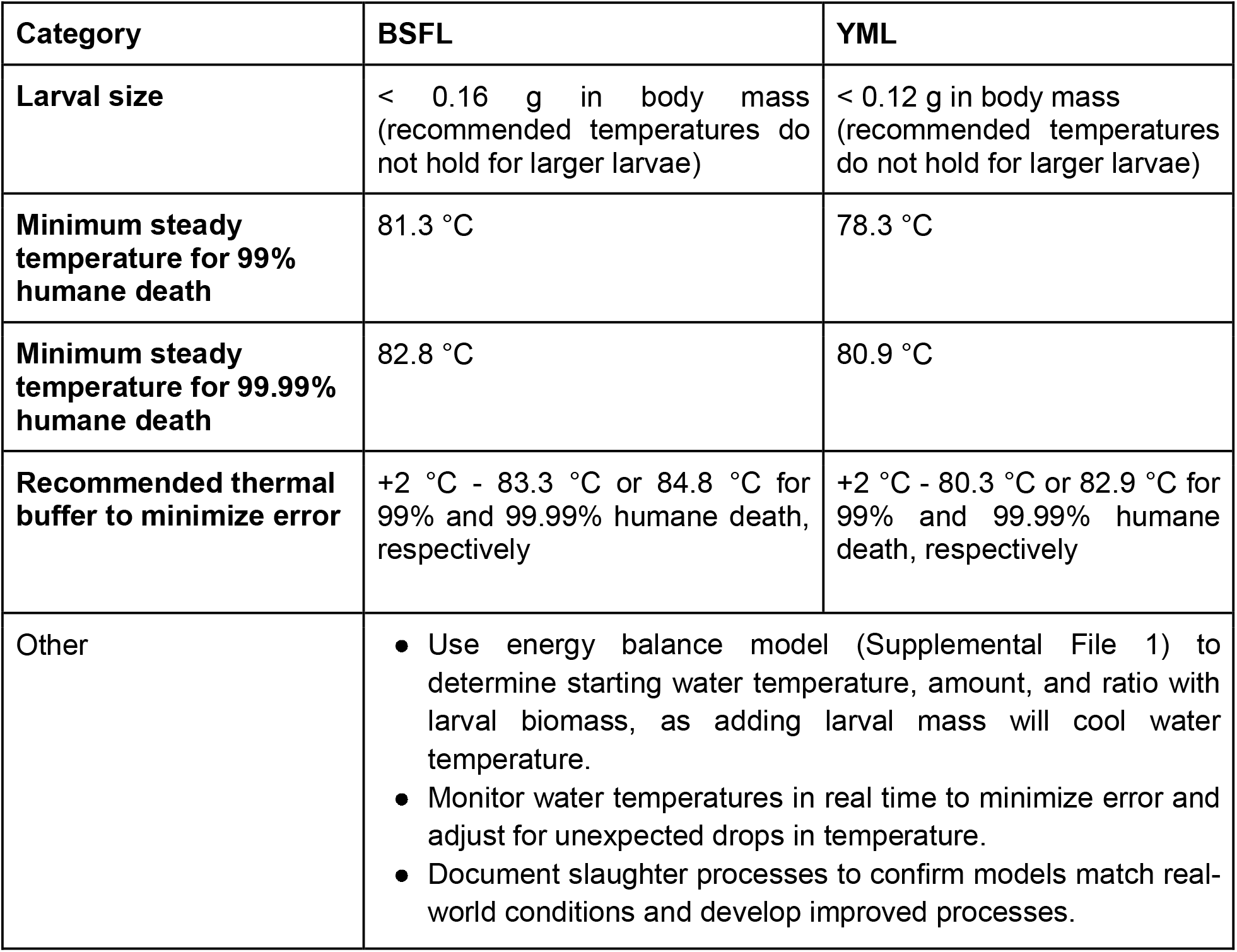
Recommendations for humane slaughter of BSFL and YML via hot water immersion.

| Category | BSFL | YML |
| --- | --- | --- |
| Larval size | < 0.16 g in body mass<br>(recommended temperatures do not hold for larger larvae) | < 0.12 g in body mass<br>(recommended temperatures do not hold for larger larvae) |
| Minimum steady temperature for 99% humane death | 81.3 °C | 78.3 °C |
| Minimum steady temperature for 99.99% humane death | 82.8 °C | 80.9 °C |
| Recommended thermal buffer to minimize error | +2 °C - 83.3 °C or 84.8 °C for 99% and 99.99% humane death, respectively | +2 °C - 80.3 °C or 82.9 °C for 99% and 99.99% humane death, respectively |
| Other | • Use energy balance model (Supplemental File 1) to determine starting water temperature, amount, and ratio with larval biomass, as adding larval mass will cool water temperature.<br>• Monitor water temperatures in real time to minimize error and adjust for unexpected drops in temperature.<br>• Document slaughter processes to confirm models match real-world conditions and develop improved processes. |  |

Overall, no consistent temperature-dependent trend was identified in the differences between the two modeling approaches (Figures 4 and 5). For BSFL, the dynamic Weibull model predicted shorter times-to-death at equivalent survival probabilities than the Baranyi-based model at 50.1 and 52.1 °C, as well as at 68.2 and 72.5 °C. For YML, shorter predicted times were observed at 42.1, 61.1, and 66.0 °C when using the dynamic Weibull equation. These differences indicate that temperature influences the relative predictive behavior of the two models.

The fitting parameters obtained from the Baranyi-based primary model were subsequently related to temperature through secondary models (Supplemental File 4). This approach allows the estimation of model parameters at temperatures that were not evaluated in the primary models due to measurement resolution, provided the temperatures still fell within the experimentally studied range in the isothermal assays. The predicted parameters can then be incorporated into the primary model (Baranyi probability distribution) to generate survival probability profiles as a function of time under different temperature conditions.

The fitting parameters μ (effective kinetic parameter) and *lag* (adaptation phase, Eq. 22) were plotted against temperatures (*T*). In the first case, a good linear experimental fitting for μ *vs*.*T* was obtained for both species (μ = 0.07919 *T* − 3.4909 ; R^2^: 0.9835; RMSE: 0.0641 for BSFL and μ = 0.0841 *T* − 3.2923 ; R^2^: 0.9911; RMSE: 0.0647 for YML), while the second plotting (*lag vs*.*T*) resulted in hyperbolic-shape curves (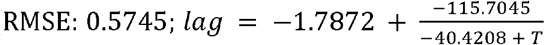 for 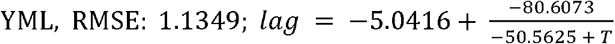 for BSFL). It is worth noting that the BSFL fittings did not include data at 50 and 56 °C, as these conditions deviated from the temperature-dependent trend observed across the other tested conditions.

Figure 6 illustrates the predicted time-to-death for survival fraction at 1% for BSFL and YML when employing the Dynamic Weibull distribution. To achieve 99% death probability with *t* ≤ *s* (humane slaughter), BSFL ≤ 0.16 g would require a slaughtering water temperature of at least 81.3 °C, while YML ≤ 0.12 g would require at least 78.3 °C (Figure 6). To achieve 99.99% death probability (*S* = 0.0001) with *t* ≤ 1 *s* for larvae of the same mass range, BSFL would require a slaughtering water temperature of at least 82.7 °C, while YML would require at least 80.9 °C. Such estimates were obtained based on Eq. 30 (always rounding up to the nearest 0.1 °C to avoid underestimating the temperature required for humane slaughter).

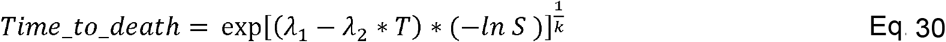

**Figure 6.**
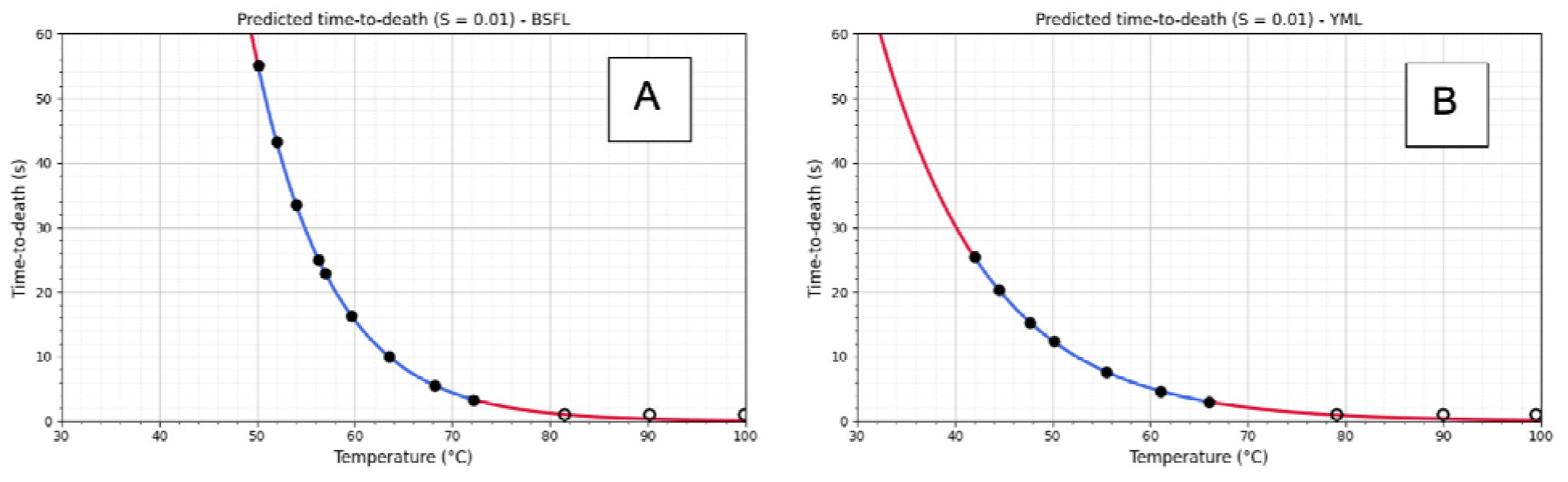
Predicted time-to-death for larvae survival fraction at 1% probability (or, 99% death probability) for A) BSFL and B) YML using the dynamic Weibull distribution that incorporates the cumulative thermal injury. The filled black circles represent estimates using the generated Weibull model for the experimental points, while the red line shows an extrapolation of the model. The empty black circles represent real data collected at those temperatures, demonstrating model fit with isothermal critical thermal maxima data collected on real animals.

Where *λ*_1_, *λ*_2_ and represent the fitting parameters when all isotherms were fitted to the model; *T* is the temperature (°C) and *S* the survival fraction. Table 1 illustrates the fitting parameters found for both insects.

As previously explained, due to the 1 s measurement resolution limit (below which no distinction can be made), it was not possible to obtain model estimation without extrapolation into the humane temperature range for both species. However, when the experimentally determined time-to-death values were plotted (empty black circles) together with the model predictions, the experimental data points collected during isothermal CTmax assays fell within the extrapolated region of the model curve (red line). This agreement between the experimental observations and the extrapolated predictions provides additional support for the applicability of the model within this temperature range.

### 3.3. Product Quality Analyses - BSFL

In order to understand how slaughtering protocols that were humane (≤1 s time-to-death) versus not humane (>1 s time-to-death) impacted product quality, we assessed three slaughtering protocols for their impact on product quality variables. These protocols were: 1) a temperature resulting in a ∼1 min time-to-death for 99% of BSFL with a 2 °C buffer (52.0 °C, for 55 s); 2) a temperature resulting in ≤1 s time-to-death for 99% of larvae with a 2 °C buffer (83.3 °C for 1 s); and 3) the highest possible temperature (99.5 -100 °C) for 1 s, which will also result in ≤1 s time-to-death for >99.99% of larvae but uses more energy.

The slaughter protocol used had no significant impacts on ash, water activity (beta regression with LRT; p > 0.05), and pH (Supplemental Material 1, Table 1, Figure 7; Welch’s ANOVA, W = 3.41, p = 0.18). Crude fat was highest using the 100 °C protocol and lowest using the 52 °C protocol (X^2^ = 13.86, p < 0.001; Sidak’s, all |z| > 2.66, all p < 0.02). Moreover, crude protein also varied by treatment (X^2^ = 12.58, p = 0.0019), as the 52 °C protocol had lower crude protein than both higher temperature protocols (52-83.3: z = -5.09, p < 0.0001; 52-100: z = -3.65, p = 0.0008; 83.3-100: z = 1.44, p = 0.39). Rancidity also differed among protocols (F = 8.15, p = 0.0195). Larvae processed at 52 °C and 83.3 °C exhibited lower rancidity than those processed at 100 °C (all p < 0.046) while no significant difference was observed between 52 and 83.3 °C (p = 0.45).

**Figure 7.**
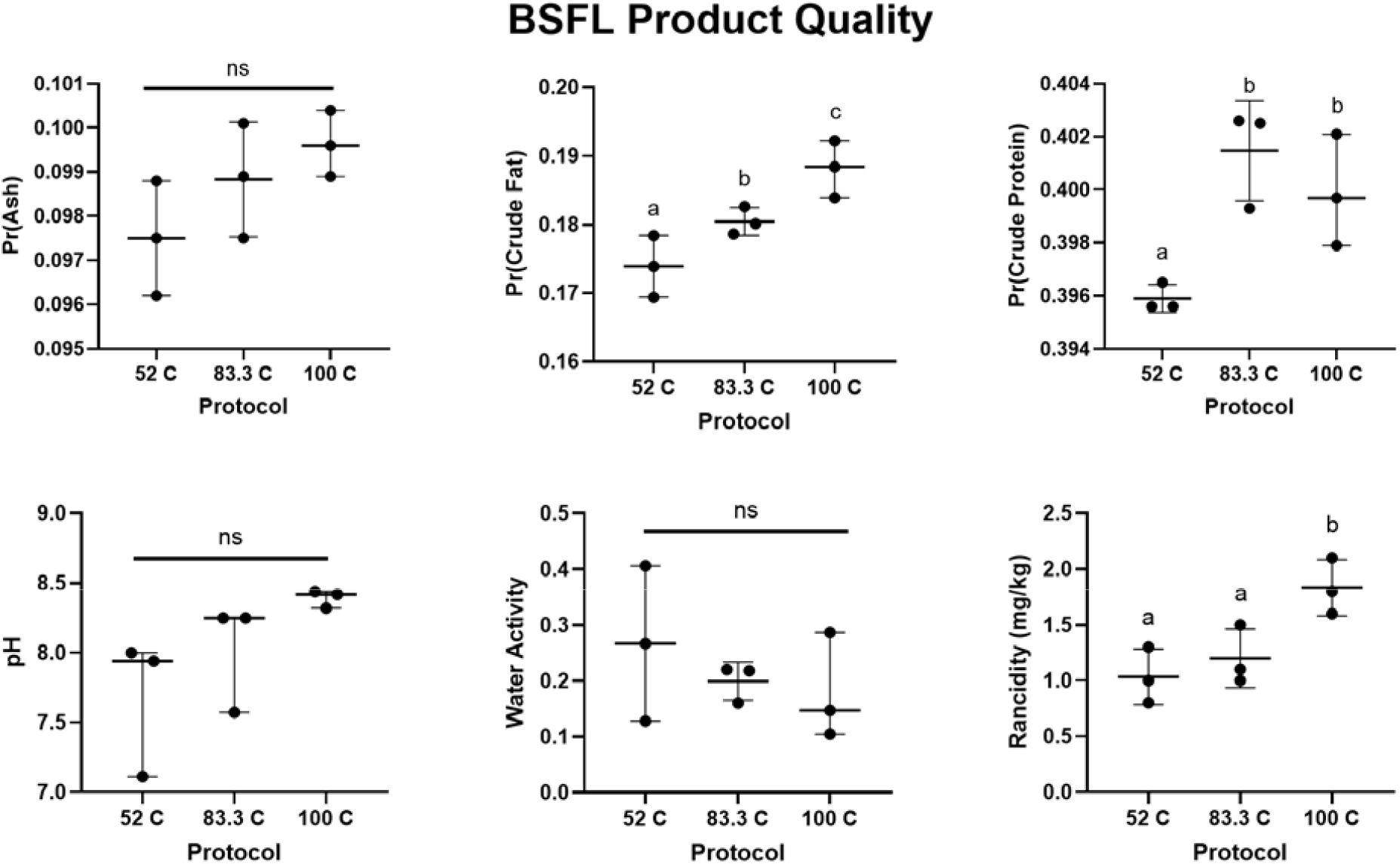
BSFL product quality measures (ash, crude fat, crude protein (Kp = 4.67), pH, water activity, and rancidity) based on slaughter protocol used. Ash, pH, and water activity did not differ significantly by protocol (p > 0.05). 100 °C performed best to maximize crude fat; 83.3 or 100 °C performed best to maximize crude protein; and 52 or 83.3 °C performed best to reduce rancidity.

Neither storage time nor protocol had an impact on flour color over 30 days of storage for BSFL flour (*L*^*\**^, *a*^*\**^, or *b*^*\**^ on day 0, 7, or 30; GLMM, all F < 12.79, p > 0.07), though a low sample size (n = 2 individual batches per protocol, with three separate flour replicates from each batch) means results should be interpreted carefully.

### 3.4. Product Quality Analyses - YML

Following the same approach used for BSFL, three slaughtering protocols were evaluated for their impact on YML product quality: 1) 47.0 °C for 55 s (temperature resulting in a ∼1 min time-to-death for 99% of larvae with a 2 °C buffer); 2) 80.3 °C for 1 s (temperature resulting in ≤1 s time-to-death for 99% of larvae with a 2 °C buffer); and the highest possible temperature (95 -100.0 °C) for 1 s.

Slaughter protocol had no significant impacts on ash (beta regression with LRT, X^2^ = 2.33, p = 0.31) and rancidity of YML (Supplemental Material 1, Table 2, Figure 8; Welch’s ANOVA, W = 1.63, p = 0.44); there was a nonsignificant trend towards higher crude protein in the 80.3 and 100 °C protocols compared to the 47 °C protocol (X^2^ = 5.58, p = 0.06). Crude fat was highest using the 80.3 and 100 °C protocols and lowest using the 47 °C protocol (X^2^ = 7.29, p = 0.026; Sidak’s, comparing 47 to 80.3 or 100 °C, both |z| > 2.83, all p < 0.014). pH was higher when YML were slaughtered using the 100 °C protocol compared to the 47 °C protocol (Kruskal-Wallis test, K-W = 7.26, p =0.0036; 100-47: p = 0.02), while pH in the 80.3 °C did not differ from either the 100 °C or 47 °C protocols (p > 0.53). Water activity was highest in the 47 °C protocol (beta regression with LRT, X^2^ = 31.03, p < 0.0001; Sidak’s, comparing 47-80.3 or 47-100 °C, both |z| > 14.37, all p < 0.0001); while the 80.3 and 100 °C protocols did not differ (p = 0.65).

**Figure 8.**
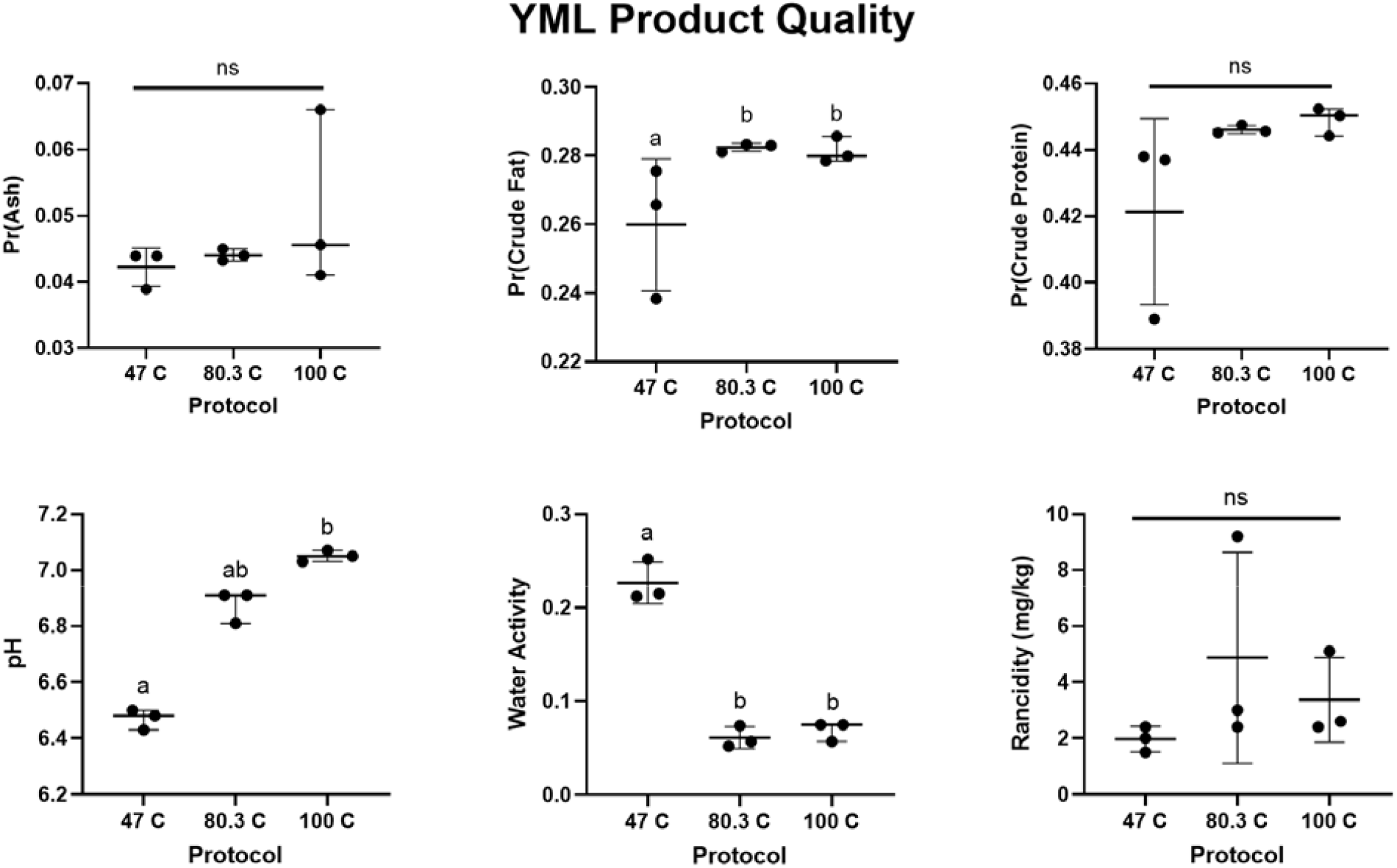
YML product quality measures (ash, crude fat, crude protein (Kp = 4.75), pH, water activity, and rancidity) based on slaughter protocol used. Ash, CP, and rancidity did not differ significantly by protocol (p > 0.05). 80.3 or 100 °C performed best to neutralize pH, preserve CF, and minimize water activity.

Storage time only impacted *a*^*\**^ values in YML flour (GLMM, F = 5.91, df = 2, 52.14, p = 0.0048; *L*^*\**^ and *b*^*\**^, F < 0.81, p > 0.45), with decreased *a*^*\**^ after 7 and 30 days of storage (Tukey’s, p < 0.028). Protocol impacted *L*^*\**^, *b*^*\**^, and *a*^*\**^ in YML flour (Table 2, all F > 8.69, p < 0.0012). 80.3 °C and 100 °C protocols increased *L*^*\**^ and *b*^*\**^ compared to 47 °C (i.e., lighter, more golden flours at higher temperatures, *L*^*\**^: p < 0.0135; *b*^*\**^, p < 0.0075), and 100 °C decreased *a*^*\**^ compared to both 80.3 °C and 47 °C (i.e., reduced red hue, p < 0.0041).

## 4. Discussion

### 4.1. Empirical conclusions

Welfare-relevant modeling remains limited, despite its critical importance for achieving humane slaughter for the trillions of insects produced annually for food and feed. In industrial insect processing, hot water immersion (often described as blanching or boiling) is commonly used as a killing step, reducing microbial loads and to inactivating endogenous enzymes responsible for browning and lipid oxidation (Caligiani *et al*., 2019). However, little attention has been paid to the exact SOPs required for humane death (≤1 s) by hot water immersion for BSFL or YML; our study aimed to assess if hot water immersion could be ‘instantaneous’. The dynamic Weibull distribution predicted a temperature of 81.3 °C for BSFL that are ≤ 0.16 g in mass, and a temperature of 78.3 °C for YML that are ≤ 0.12 g, would be necessary to kill 99% of larvae within 1 s (i.e., humanely). Further, using killing SOPs based on these temperatures had no negative impacts on product quality or color and, in some cases, even benefited product quality and color compared to prolonged killing at lower water temperatures.

Heat death in insects may be caused by central nervous system (CNS) failure, cellular damage, protein denaturation, and/or changes in membrane fluidity. A study in fruit flies showed that spreading depolarization in the CNS reliably correlates with heat coma (e.g., the ‘paralysis’ measure taken to indicate onset of insensibility and plausible death in our study) though CNS failure may not necessarily be the proximate cause of death (Jørgensen *et al*., 2020). Our study design is different from most CTmax studies conducted with insects, which often use a ramping assay to find the ‘lower bound’ CTmax temperature (Overgaard *et al*., 2012); our data were instead generated using isothermal conditions and assessing time-to-death for each individual larva, and therefore should not be compared with ramping assay results. However, our study design is generalizable to the context we were modeling, where insects are dropped in already-heated water. We found that survival ratios decreased with increased processing time at the lower water temperatures in the Weibull probability distribution models for both species; this finding is consistent with previous work demonstrating a decrease in the survival ratio of maize weevil adults with increasing processing time during hot-air disinfestation at temperatures ranging from 46 to 62 °C (Parisotto *et al*., 2023).

The Weibull probability distribution models displayed a common curve behavior for both species, with lower temperatures exhibiting a pronounced shoulder at the beginning of the curves (i.e., at short processing times), which may indicate the animals have a longer period of initial tolerance at these temperatures. This was followed by a rapid exponential decline in the surviving population until a final plateau was reached, corresponding to complete slaughtering. Despite the curves having the same sigmoidal shape, it is worth noting that YML presented a substantially shorter initial shoulder than BSFL at lower temperatures, which suggests a greater thermal resistance for the BSFL larvae. Similar kinetic patterns have been reported for maize weevils, with higher temperatures resulting in shorter processing times and a reduced initial shoulder in the sigmoidal survival curve (Parisotto *et al*., 2023).

Additionally, to the best of our knowledge, no previous studies have directly compared the thermal tolerances of YML and BSFL under the same CTmax conditions. However, studies of these insects in farmed conditions also suggest greater tolerance in BSFL: crates of 10,000 BSFL are known to generate heat up to 45.6 °C without significant lethality (Li *et al*., 2023) while, in YML, previous research has reported that in-crate temperatures of >40 °C (with a high of 43.9 °C) can be lethal (Deruytter *et al*., 2022).

All isothermal CTmax data was taken using individual larvae dropped onto the surface of the hot water; these larvae naturally floated on the water surface in most cases and this likely reduced the rate at which their body increased in temperature (and thus increased time-to-death) compared to if they had been totally immersed in the water. However, we believe this replicates real-world conditions, where many larvae also float to the top of the water when being killed (even if some others may be forced below the water surface due to the volume of larval biomass above them). We have thus conservatively modeled the temperature required to kill 99% of larvae in less than one second by assuming all the animals are ‘surface-sitting’ larvae and not totally immersed. In other words, whether producers intentionally submerge their larvae completely or not, our model will not have incorrectly predicted a lower temperature than what is actually required to achieve a humane death for 99% of them.

We also found that producers do not need to choose between ethical and economical killing when blanching. One or both of the SOPs we used that resulted in ≤1 s time-to-death performed best to maximize crude fat and protein, as well as reduce rancidity, in BSFL. In YML these SOPs performed best to neutralize pH, maximize crude fat, and minimize water activity, and may also maximize crude protein. These results are not necessarily surprising: thermal enzyme-inactivating methods, particularly blanching, consistently result in superior protein stability: In *H. illucens* larvae, blanching (typically 40–60 s at ∼100 °C) markedly reduced indicators of lipid oxidation and protein-related browning compared to non-thermal or slow thermal killing methods (Leni *et al*., 2019). Similarly, a study by Gomes da Costa Lima *et al*., (2026) found that boiling yellow mealworm at 100 °C (for 10 minutes) resulted in a combination of high crude protein and lighter-colored final flours, which was not found with many other slaughter methods. Similarly, our results showed that the higher temperature blanching conditions we recommend as humane resulted in increased YML flour lightness and provided a more golden color profile (higher *L*^*\**^ and *b*^*\**^) that was very stable over 30 days of storage, while BSFL flours were unaffected by protocol used (though, perhaps due to low sample size).

The impact of the slaughter method on total lipid content is often modest (e.g., the small difference in crude fat between blanching and freezing in Martins da Silva *et al*., 2024). Our study showed similarly modest impacts on crude fat in both species under humane, high-temperature SOPs compared to the lower temperature, more prolonged killing SOP. While we did not test this, the distribution of lipid classes may be impacted more significantly by killing method: BSF prepupae killed by freezing saw a dramatic increase in free fatty acids as a result of extensive lipolysis, while blanching inactivated endogenous lipases and preserved triacylglycerol-rich lipid fractions (Caligiani *et al*., 2019). Rapid thermal treatments, like our recommended humane protocols, may thus stabilize lipid fractions by inactivating hydrolytic enzymes. Blanching also reduces lipid oxidation rates during storage compared to methods like blending, freezing, vacuum, or CO_2_ treatments (Zhen *et al*., 2020), further demonstrating that ethical and economical practice may be aligned when using hot water immersion for killing.

### 4.2. Recommendations for production

Keeping water above a temperature of 81.3 °C for BSFL that are ≤ 0.16 g in mass, and above a temperature of 78.3 °C for YML that are ≤ 0.12 g, is recommended to kill 99% of larvae within 1 s (i.e., humanely). Importantly, starting water temperature will need to be higher than these temperatures, as adding a large amount of larval biomass to the hot water will drop the water temperature to a point below the recommendations. Producers are thus advised to use the energy balance model (Supplementary File 1) to determine the water starting temperature and volume, and larvae:water ratio, needed to achieve humane slaughter with hot water immersion in their facilities. Producers should also monitor water temperatures in their facility when engaging in slaughter to document the use of appropriate slaughter SOPs and confirm our models match their real-world conditions; this can also aid producers looking to differentiate their products from others via humane labeling by ensuring they have a record of using best practices (Barrett and Adcock, 2023).

Producers may consider using a safety margin of at least 2 °C (i.e., 83.3 °C for BSFL and 80.3 °C for YML), as we did in our product quality tests, to err on the side of caution in case larval mass or water volume/temperature are not on target; this would reduce the likelihood of accidental inhumane slaughter for thousands to millions of animals per batch. Even just in our small laboratory setup (145 g of larvae, 8 L of hot water), water temperature dropped by as much as 0.7 °C as a result of adding the room temperature basket and larval biomass. Water temperature should always be stable at the desired start temperature before introducing larvae.

Two limitations should be considered for producers adopting these temperatures as guidance for humane killing in their production facility. First, our pilot data suggests that increased body size led to longer time-to-death in both species at many of the tested temperatures (Supplementary Materials, Figure S1 and S2). This means that our recommended water temperatures should only be used with animals within or below the size ranges we tested, as our recommendations may underestimate the temperature needed to guarantee humane death in BSFL and YML that are larger than our tested sizes (< 0.16 g for BSFL; < 0.12 g for YML). However, our pilot data also suggest the effect of body size is more pronounced at lower temperatures that are unlikely to be instant. For instance, death time increased by 4.8 s per 0.1 g of body mass in BSFL at 56 °C but only 1.8 s per 0.1 g of body mass at 72 °C. Therefore, higher temperatures at or above our recommended SOPs will have reduced error associated with body mass variation compared to lower, non-recommended temperatures. Still, further research should assess the impact of body size on death time more concretely for producers that may be slaughtering much larger larvae; there may be very large larval body sizes for which no hot water temperature will prove ‘instant’/humane.

Second, these recommendations are designed to be used only with animals that have been separated from substrate prior to hot water immersion. Depopulation events may require producers to kill large numbers of larvae that are still embedded in substrate. Substrate characteristics may influence the time-to-death via hot water immersion; for instance, a very wet substrate or one with significant fungal growth may not break apart easily in water. This will cause the larvae to heat more slowly, as they are prevented from directly contacting the water due to the insulating substrate around them. Further research would be needed to assess how substrate characteristics may affect humane killing via hot water immersion for depopulation events.

Recommendations for producers interested in the humane slaughter of BSFL and YML via hot water immersion are summarized in Table 3.

## 5. Conclusion

The present study addresses an important knowledge gap by providing a comparative assessment of BSFL and YML thermal survival responses to hot water treatments and enabling the prediction of the insect survival probabilities under different thermal conditions. 99% of BSFL ≤ 0.16 g and YML ≤ 0.12 g in mass can be humanely slaughtered in one second or less when water temperatures are maintained above 81.3 °C and 78.3 °C, respectively. Both the dynamic Weibull distribution and the Baranyi model reasonably described the survival curve of BSFL and YML, as confirmed by the low RMSE values and agreement between the predicted and observed time-to-death in the experimental Ctmax assays for larvae survival fraction at 1% probability. The energy balance equation is a simple tool for producers looking to develop humane SOPs in their facility, as it can help to determine key process parameters for humane slaughtering while requiring minimum input. Documenting water temperatures throughout slaughter, larval biomass, and water volume can help producers maintain high-quality records of humane slaughter in their facility. Finally, our product quality data show that humane blanching SOPs have neutral or beneficial effects on product quality, showing that humane and economical slaughter practices are possible for larvae of specific body sizes when slaughtered via hot water immersion. Further research on other species, the effect of body size on time-to-death, and the effects of substrate on time-to-death when using hot water immersion for depopulation events may further elucidate the parameters that generate humane larval killing for a variety of insect production systems and killing needs.

## Acknowledgments

Thanks to Alexander Glica for his help and support with this work. Narmeen Hashim, Charlie Schmidt, Elijah Persson-Gordon, and Talia Bair who helped prepare larvae for the product quality SOP tests; we are very grateful for their work.

## Supplemental Materials

**Supplemental Figure 1.**
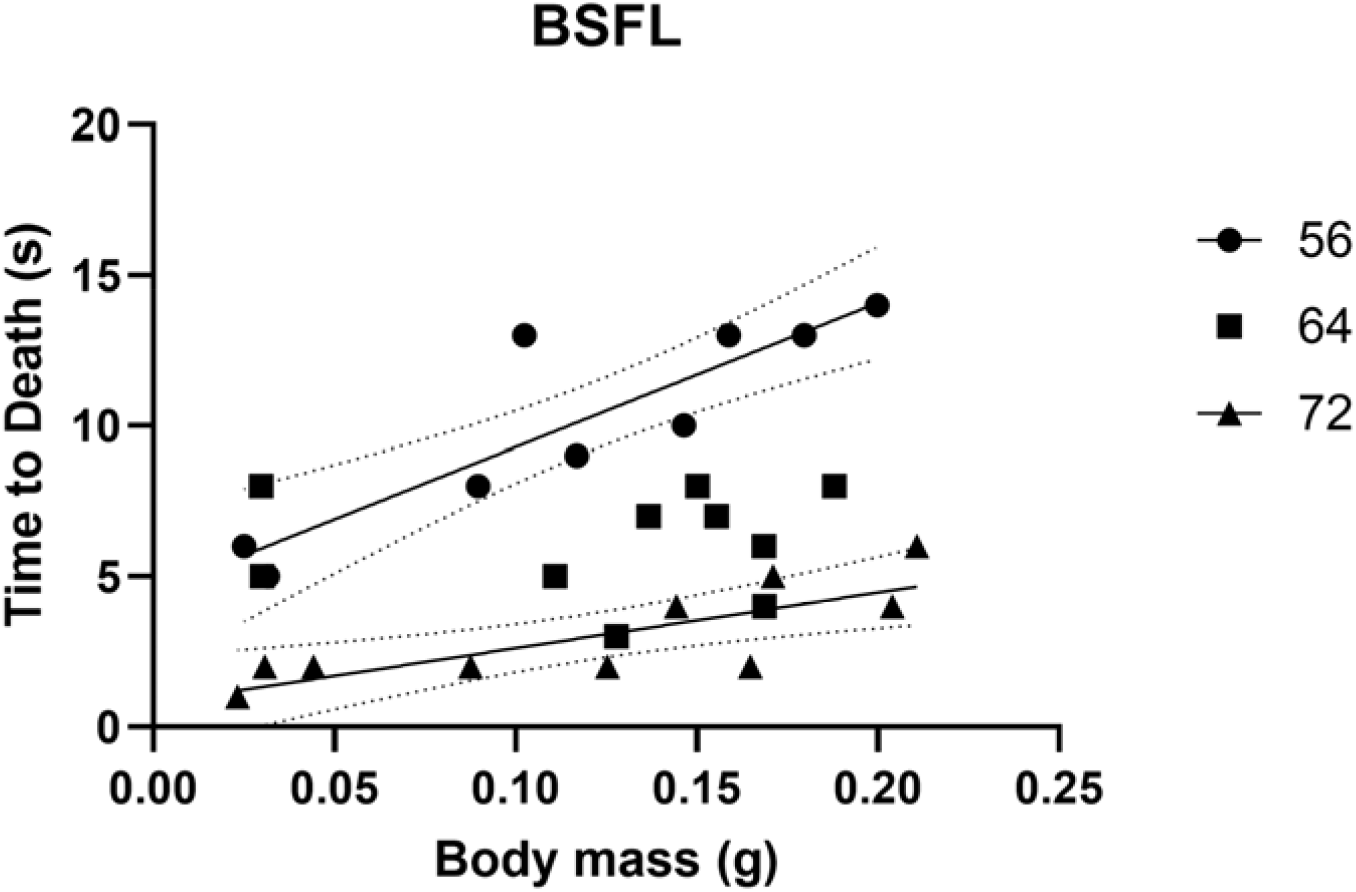
BSFL body size predicted time to death at 56 and 72 °**C, but not 64** °**C**. Smaller body sizes resulted in faster death times for larvae placed in 56 and 72 °C water (p < 0.0059) but not 64 C water (p = 0.90). For 56 °C: death time (s) = 47.93^*^ mass (g) + 4.50 (F = 32.56, df = 8, R^2^ = 0.80). For 72 C: death time (s) = 18.45^*^ mass (g) + 0.77 (F = 13.81, df = 8, R^2^ = 0.63). Solid lines indicate significant relationships with dotted 95% CI.

**Supplemental Figure 2.**
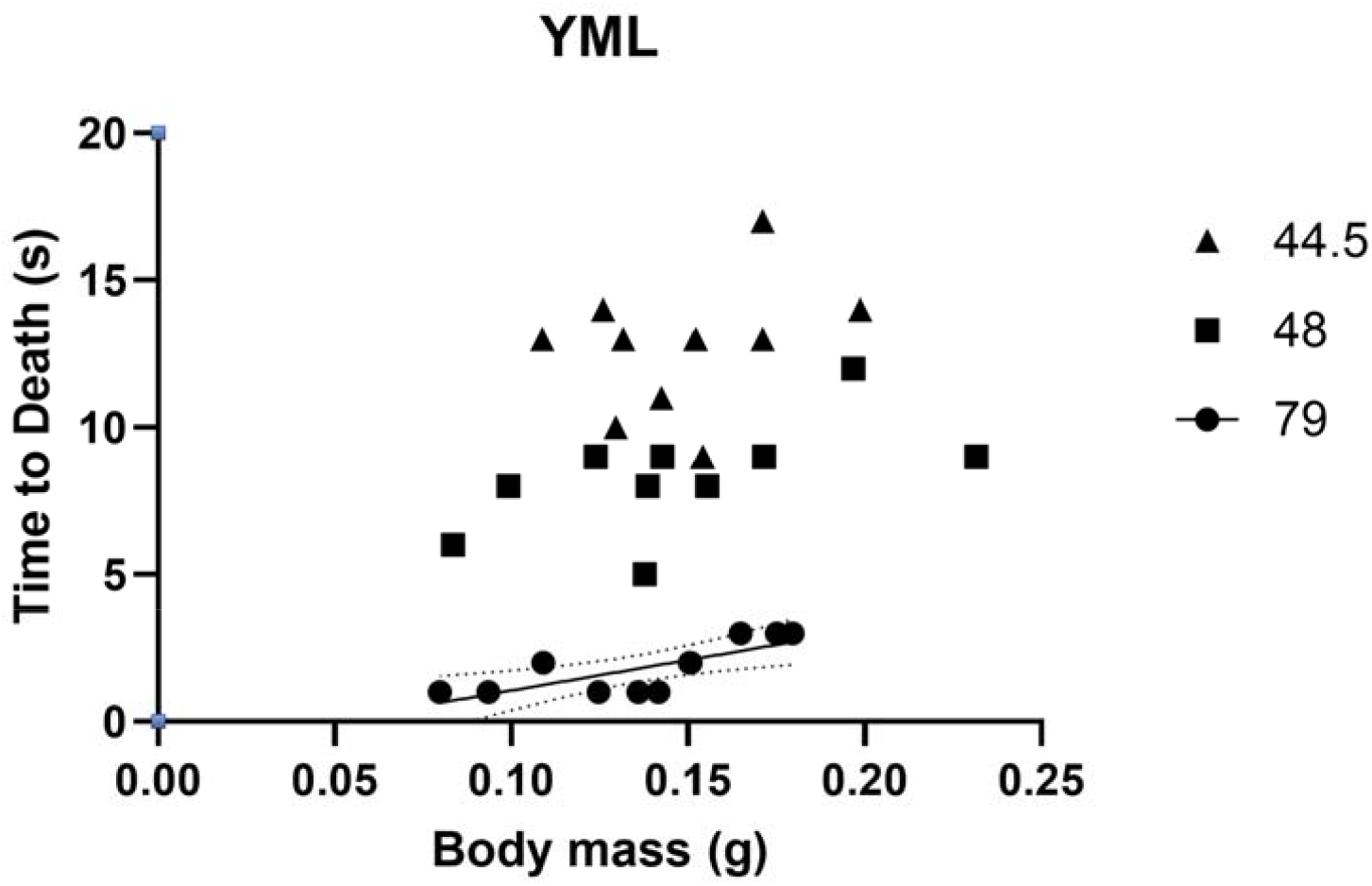
YML body size predicted time to death at 79 °**C, but not 44.5 or 48** °**C**. Smaller body sizes resulted in faster death times for larvae placed in 79 °C water (p < 0.0098) but not 44.5 (p = 0.34) or 48 °C water (p = 0.079). For 48 °C: death time (s) = 24.97^*^mass + 4.60. For 79 °C: death time (s) = 20.81^*^ mass (g) - 1.02 (F = 11.36, df = 8, R^2^ = 0.59). Solid lines indicate significant relationships with dotted 95% CI; dashed lines indicate non-significant trend.

**Supplemental Table 1.** Mean ± SD or median product quality measures (n = 3/protocol) for BSFL based on slaughter protocol used, reported on DM basis.

| Process Temp (time) | Ash (% DM) | Crude Fat (% DM) | Crude Protein (% DM; 4.67 N:P) | Crude Protein (% DM; 6.25 N:P) | pH<sup>1</sup> | Water activity (aw) | Rancidity (mg/kg) |
| --- | --- | --- | --- | --- | --- | --- | --- |
| 52.0 °C (55 sec) | 9.75±0.13<sup>a</sup> | 17.4±0.45<sup>a</sup> | 39.59±0.05<sup>a</sup> | 52.98±0.07<sup>a</sup> | 7.94<sup>a</sup> | 0.26±0.14<sup>a</sup> | 1.03±0.25<sup>a</sup> |
| 83.3 °C (1 sec) | 9.88±0.13<sup>a</sup> | 18.0±0.20<sup>b</sup> | 39.99±0.18<sup>b</sup> | 53.52±0.25<sup>b</sup> | 8.25<sup>a</sup> | 0.21±0.03<sup>a</sup> | 1.20±0.26<sup>a</sup> |
| 100.0 °C (1 sec) | 9.96±0.08<sup>a</sup> | 18.8±0.42<sup>c</sup> | 40.14±0.21<sup>b</sup> | 53.73±0.25<sup>b</sup> | 8.42<sup>a</sup> | 0.18±0.10<sup>a</sup> | 1.83±0.25<sup>b</sup> |

<sup>1</sup>Non-normal; reported as median
DM: Dry mass
<sup>a</sup> Letters indicate statistically significant differences among treatments in that column

**Supplemental Table 2.** Mean ± SD or median product quality measures (n = 3/protocol) for YML based on slaughter protocol used, reported on DM basis.

| Process Temp (time) | Ash (% DM)<sup>1</sup> | Crude Fat (% DM) | Crude Protein (% DM; 4.75 N:P)<sup>1</sup> | Crude Protein (% DM; 6.25 N:P)<sup>1</sup> | pH<sup>1</sup> | Water activity (aw) | Rancidity (mg/kg)<sup>1</sup> |
| --- | --- | --- | --- | --- | --- | --- | --- |
| 47.0 °C (55 sec) | 4.39<sup>a</sup> | 26.0±1.92<sup>a</sup> | 43.70<sup>a</sup> | 57.50<sup>a</sup> | 6.48<sup>a</sup> | 0.23±0.02<sup>a</sup> | 2.00<sup>a</sup> |
| 80.3 °C (1 sec) | 4.40<sup>a</sup> | 28.2±0.12<sup>b</sup> | 44.56<sup>a</sup> | 58.63<sup>a</sup> | 6.91<sup>ab</sup> | 0.06±0.01<sup>b</sup> | 3.00<sup>a</sup> |
| 100.0 °C (1 sec) | 4.56<sup>a</sup> | 28.1±0.38<sup>b</sup> | 45.03<sup>a</sup> | 59.25<sup>a</sup> | 7.05<sup>b</sup> | 0.07±0.01<sup>b</sup> | 2.60<sup>a</sup> |

<sup>1</sup>Non-normal data are reported as medians
<sup>a</sup> Letters indicate statistically significant differences among treatments in that column

**Supplemental Table 3.** Root mean square error (RMSE) comparing the Weibull survival function incorporation of cumulative thermal injury (Dynamic Weibull) and the Weibull-Baranyi model (Baranyi) with the traditional Weibull survival function for black soldier fly (BSFL) and yellow mealworm (YML) larvae.

| RMSE - BSFL |  |  |
| --- | --- | --- |
| Temperature (°C) | Dynamic Weibull | Baranyi |
| 50.1 | 0.1826 | 0.0142 |
| 52.1 | 0.1721 | 0.0101 |
| 54.0 | 0.0676 | 0.0299 |
| 55.7 | 0.2529 | 0.0290 |
| 57.5 | 0.1317 | 0.0522 |
| 59.5 | 0.0842 | 0.0132 |
| 63.4 | 0.0832 | 0.0471 |
| 68.2 | 0.0316 | 0.0369 |
| 72.5 | 0.1689 | 0.0224 |
| RMSE - YML |  |  |
| Temperature (°C) | Dynamic Weibull | Baranyi |
| 42.1 | 0.0944 | 0.0080 |
| 44.6 | 0.0058 | 0.0211 |
| 47.8 | 0.0888 | 0.0200 |
| 50.1 | 0.0738 | 0.0097 |
| 55.4 | 0.1626 | 0.0712 |
| 61.1 | 0.1512 | 0.0499 |
| 66.0 | 0.0311 | 0.0225 |

## Notes

**Conflicts of Interest:** MB reports a relationship with Rethink Priorities (consultant) that ended in 2023. Rethink Priorities had no oversight of the methodology, data collection or analysis, writing, or decision to publish this work. MB is Director of the Insect Welfare Research Society (unpaid); on the publications and open science committee of the Royal Entomological Society (unpaid); and lead editor at *Approaches in Entomology* and associate editor at *Journal of Insects as Food and Feed* (unpaid). SOD is an Assistant Development Officer at the International Society for Applied Ethology (unpaid).

### Competing Interest Statement

MB reports a relationship with Rethink Priorities (consultant) that ended in 2023. Rethink Priorities had no oversight of the methodology, data collection or analysis, writing, or decision to publish this work. MB is Director of the Insect Welfare Research Society (unpaid); on the publications and open science committee of the Royal Entomological Society (unpaid); and lead editor at Approaches in Entomology and associate editor at Journal of Insects as Food and Feed (unpaid). SOD is an Assistant Development Officer at the International Society for Applied Ethology (unpaid).

