## Supplemental File 4 for "Mathematical modelling of thermal death kinetics to describe humane slaughter procedures via hot water immersion in black soldier fly and yellow mealworm larvae"

**Supplementary File 4**

Figure S4.1 – Temperature dependence of Weibull–Baranyi secondary model parameters for yellow mealworm. The red points were experimental data, however, not considered for the fitting since it did not follow the trend.

Figure S4.2 – Temperature dependence of Weibull–Baranyi secondary model parameters for black soldier fly. The red points were experimental data, however, not considered for the fitting since it did not follow the trend.
