## Supplemental File 3 for "Mathematical modelling of thermal death kinetics to describe humane slaughter procedures via hot water immersion in black soldier fly and yellow mealworm larvae"

**Supplementary File 3**

**Calculation of heat capacity**

The heat capacity of each larva ($C_{p,L}$), used in the energy balance, was estimated based on its proximate composition (Table S3.1), obtained from selected studies in the literature (Supplemental File 2). The compositional data were then incorporated into the equation proposed by Heldman and Singh (1981) (Eq. 1):

Table S3.1 – Centesimal composition of BSFL and YML based on the average of values found in the literature.

| Component | BSFL | YML |
| --- | --- | --- |
| Protein (%, d.b.) | 39.96 | 46.78 |
| Fat (%, d.b.) | 28.41 | 30.19 |
| Carbohydrate (%, d.b.) | 20.97 | 20.56 |
| Ash (%, d.b.) | 10.47 | 4.75 |
| Moisture (%, w.b.) | 55.77 | 65.00 |

d.b.: dry basis while

w.b.: wet basis

$C_{pL}=\left( 1.424 X_{C}+1.549 X_{P}+1.675 X_{f}+0.837 X_{a} \right)*\left( 1- U_{w} \right)+4.187*U_{w}$	(Eq. 1)

Where $X$ is the mass fraction (g component/g dry solids) and the subscripts represent carbohydrate ($C$), protein ($P$), fat ($f$), and ash ($a$). $U_{w}$ represents the moisture content in wet basis (g water/g total).

As a practical example, for BSFL larvae, if the moisture content is 55.77%, then the heat capacity can be calculated as follows:

$C_{pL}=\left( 1.424*0.2097+1.549* 0.3996+1.675*0.2841+0.837*0.1047 \right)*\left( 1- 0.5577 \right)+4.187*0.5577$

$C_{pL, BSFL}=$3.0 (kJ ⋅ kg^-1^ ⋅°C^-1^)

In the same way, for YML, if the moisture content is 65%, then the heat capacity can be calculated as follows:

$C_{pL}=\left( 1.424*0.2056+1.549* 0.4678+1.675*0.3019+0.837*0.0475 \right)*\left( 1- 0.65 \right)+4.187* 0.65$

$C_{pL, YML}=$ 3.27 (kJ /kg⋅°C)
